# DAF-18/PTEN prevents oocyte wastage in spermless *C. elegans* hermaphrodites by blocking spermatheca neck dilation through RHO-1/RhoA disinhibition

**DOI:** 10.1101/2025.09.16.676617

**Authors:** Jichao Deng, Armi M. Chaudhari, Olivier Gagné, Vincent Roy, Benjamin Dufour, Pier-Olivier Martel, Martin J. Simard, Patrick Narbonne

## Abstract

Loss of the Phosphatase and TENsin homolog PTEN drives cancer in humans. In *C. elegans* hermaphrodites that lack sperm, the PTEN ortholog DAF-18 is required for oocyte quiescence and their proximal accumulation, along with the concomitant homeostatic downregulation of germline stem cell proliferation. As such, spermless *daf-18(ø)* mutants undergo full-scale oogenesis and ovulation in a futile manner, wasting all produced oocytes. When oocyte laying is prevented in these mutants, oocytes hyperaccumulate to form differentiated benign tumors. Here, we show that *daf-18* expression in the spermatheca neck myoepithelium is sufficient to induce oocyte arrest and homeostatic downregulation of germline stem cell proliferation in spermless hermaphrodites, altogether prohibiting tumorigenesis. We demonstrate that DAF-18 promotes spermatheca neck contractility through limiting PIP_3_ levels and AKT activity, preventing the latter from inactivating RHO-1/RhoA by direct phosphorylation at a conserved site. Loss of PTEN may therefore promote benign tumorigenesis in humans through similar cell non-autonomous effects.

## Introduction

The Phosphatase and TENsin homolog PTEN is a dual-specificity phosphatase best known for dephosphorylating phosphatidylinositol (3,4,5)-triphosphate (PIP_3_) into phosphatidylinositol (4,5)-diphosphate (PIP_2_) (Song et al., 2012). Decreased PIP_3_ levels in turn reduce AK strain transforming (AKT) activation and mammalian target of rapamycin (mTOR) signaling to downregulate protein synthesis, and hence, cell growth and proliferation (Kriplani et al., 2015; Manning and Cantley, 2007). AKT inactivation also mobilizes forkhead box O (FOXO) transcription factors, which alter gene expression to establish and consolidate a state of cellular dormancy (Burgering and Medema, 2003). Consequently, PTEN is an important tumor suppressor, the loss of which is a driver in multiple types of cancers (Solimini et al., 2012; Song et al., 2012). PTEN is so paramount in preventing overproliferation that even subtle reductions in activity or protein levels through hemizygosity promote the development of multiple benign tumors in several tissues, through rare heritable conditions known as PTEN hamartoma tumor syndromes (PHTS) (Lee et al., 2018; Liaw et al., 1997; Marsh et al., 1997). While there is overwhelming evidence supporting cell autonomous tumor suppressor roles for PTEN across cell types and organisms (Chen et al., 2018; Knobbe et al., 2008), here we report a situation where the *C. elegans* PTEN ortholog *daf-18* acts exclusively cell non-autonomously to prevent benign tumorigenesis.

In *C. elegans*, loss-of-function mutations in the gene coding for the PTEN ortholog *daf-18* were initially identified through their <u>da</u>uer formation defective (Daf-d) phenotype (Ogg and Ruvkun, 1998). When young *C. elegans* larvae encounter harsh environmental conditions leading to the downregulation of insulin/IGF-1 signaling (IIS), DAF-18 is required to activate the forkhead transcription factor DAF-16/FOXO, which in turn, induces dauer entry (Ogg et al., 1997; Ogg and Ruvkun, 1998). Crowding can nonetheless promote dauer formation independently from DAF-18 and DAF-16, via the downregulation of transforming growth factor beta (TGF-β) signaling (Narbonne and Roy, 2006a; Ogg et al., 1997). In this case, *daf-18(ø)* dauers contain extra germline stem cells (GSCs) and gonad blast cells, while *daf-16(ø)* dauers do not show such defects (Narbonne and Roy, 2006b; Tenen and Greenwald, 2019). DAF-18 can therefore block cell proliferation independently from DAF-16 and in this context, it was intriguingly shown to act cell non-autonomously from the somatic gonad to promote both GSC and gonad blast quiescence (Tenen and Greenwald, 2019). The mechanism for this DAF-16-independent and cell non-autonomous tumor suppressive role of DAF-18 has however remained obscure.

When *daf-18(*ø) larvae are grown under optimal conditions, they surprisingly do not show any overt defects, with the exception of being short-lived as adults (Mihaylova et al., 1999). DAF-18 is otherwise implicated in oocyte maturation though defects are only apparent under specific conditions (Brisbin et al., 2009; Suzuki and Han, 2006). A dramatic defect of *daf-18(*ø) mutants is however revealed in older sperm-depleted hermaphrodites, or germline feminized (*fog*) *“female”* mutants. Sperm is required to release oocytes from diakinesis and promote ovulation, and thus arrested oocytes accumulate in the proximal gonad of spermless hermaphrodites (Miller et al., 2001). This induces a homeostatic signal that inhibits GSC proliferation and differentiation to pause oocyte production; the oocyte backlog being only released upon sperm reception (Cinquin et al., 2016; Morgan et al., 2010; Narbonne et al., 2015). In spermless *fog-1; daf-18(*ø) double mutants however, oocytes do not arrest in meiotic prophase, are ovulated, and undergo endomitosis (McCarter et al., 1999; Narbonne et al., 2015, 2017). Oocytes are regarded as precious cells that uniquely possess the capacity to propagate the species and often require the sacrifice of many other cell’s cytoplasm to dramatically expand in size (Gumienny et al., 1999). Despite this, *fog-1; daf-18(*ø) double mutants sadly waste them all since their GSCs keep proliferating as in sperm-bearing hermaphrodites to sustain, in vain, a full-blown oogenic program. When oocyte activation and laying is blocked, the phenotype becomes even more dramatic as homeostatic signaling defects cause oocytes to hyperaccumulate and form differentiated benign tumors (Narbonne et al., 2017). The formation of benign differentiated tumors in multiple tissues is a feature of PTEN hemizygosity in humans that may therefore arise because of defective homeostatic signaling (Valet and Narbonne, 2022).

Here we investigated how DAF-18 prevents spermless *C. elegans* hermaphrodites from wasting their precious oocytes. We found that DAF-18 functions cell non-autonomously in the animal’s spermatheca (Sp) neck myoepithelium to promote contractility and prevent the spontaneous entry of oocytes, ensuring their meiotic arrest and accumulation. This permits homeostatic signaling to the GSCs and prevents their unnecessary proliferation. We provide evidence that DAF-18 does not promote Sp neck contractility through its enzymatic product, PIP_2_, which is implicated in PLC/ITR-mediated Ca^2+^ signaling. Rather, we show that DAF-18 negates Sp neck dilation by reducing the levels of its substrate, PIP_3_, and downregulating AKT-1/2. AKT-1/2, in turn, promote Sp neck dilation by directly phosphorylating the RhoA ortholog RHO-1 on a highly conserved site to negate its myosin stimulating activity. As such, our results reveal how DAF-18 non-autonomously prevents unnecessary stem cell proliferation and the overproduction and wastage of their terminally differentiated progeny, or in other words benign tumorigenesis, in one tissue by affecting contractility signaling via AKT and RhoA in a neighboring tissue.

## Results

### Oocytes arrest independently from germline DAF-18

In control hermaphrodites, endogenously tagged DAF-18::mNG is ubiquitously expressed and is present in small quantities at the membrane of mature oocytes (Figure 1A) (Brisbin et al., 2009; Masse et al., 2005; Suzuki and Han, 2006). In spermless *fog-1* mutants however, we observed markedly higher levels of DAF-18::mNG at the membrane of arrested oocytes (Figures 1A and 1B). Given this expression pattern and DAF-18’s importance for downregulating germline MPK-1 and preventing oocyte maturation (Brisbin et al., 2009; Suzuki and Han, 2006), we hypothesized that germline DAF-18 was required for oocyte quiescence in the absence of sperm. We therefore introduced a germline-specific *daf-18*-rescuing transgene (Dickinson et al., 2013; Frokjaer-Jensen et al., 2008), henceforth *germline::DAF-18*, in the genome of *fog-1; daf-18(*_ø_) mutants. Since *daf-18(*ø) mutant larvae are maternally rescued for dauer entry (Gil et al., 1999), the larval progeny of *germline::DAF-18* animals may carry diluted somatic DAF-18 activity in addition to robust germline expression (Figure S1A). Unexpectedly, these transgenic animals continued to waste their oocytes like their *fog-1; daf-18(*ø) parents (Figures 1C and 1D). Moreover, GSC proliferation (as inferred from the GSC mitotic index [MI]) (Crittenden et al., 2006; Narbonne et al., 2015, 2017; Robinson-Thiewes et al., 2021) stayed elevated in *fog-1; daf-18(*ø*); germline::DAF-18* adults, as in wild-type and *fog-1; daf-18(*ø) controls (Figures 1E and 1F). These results suggested that the presence of DAF-18 in the germline cannot compensate for the ovulation and GSC proliferation defects of *daf-18(*ø) mutants.

**Figure 1.**
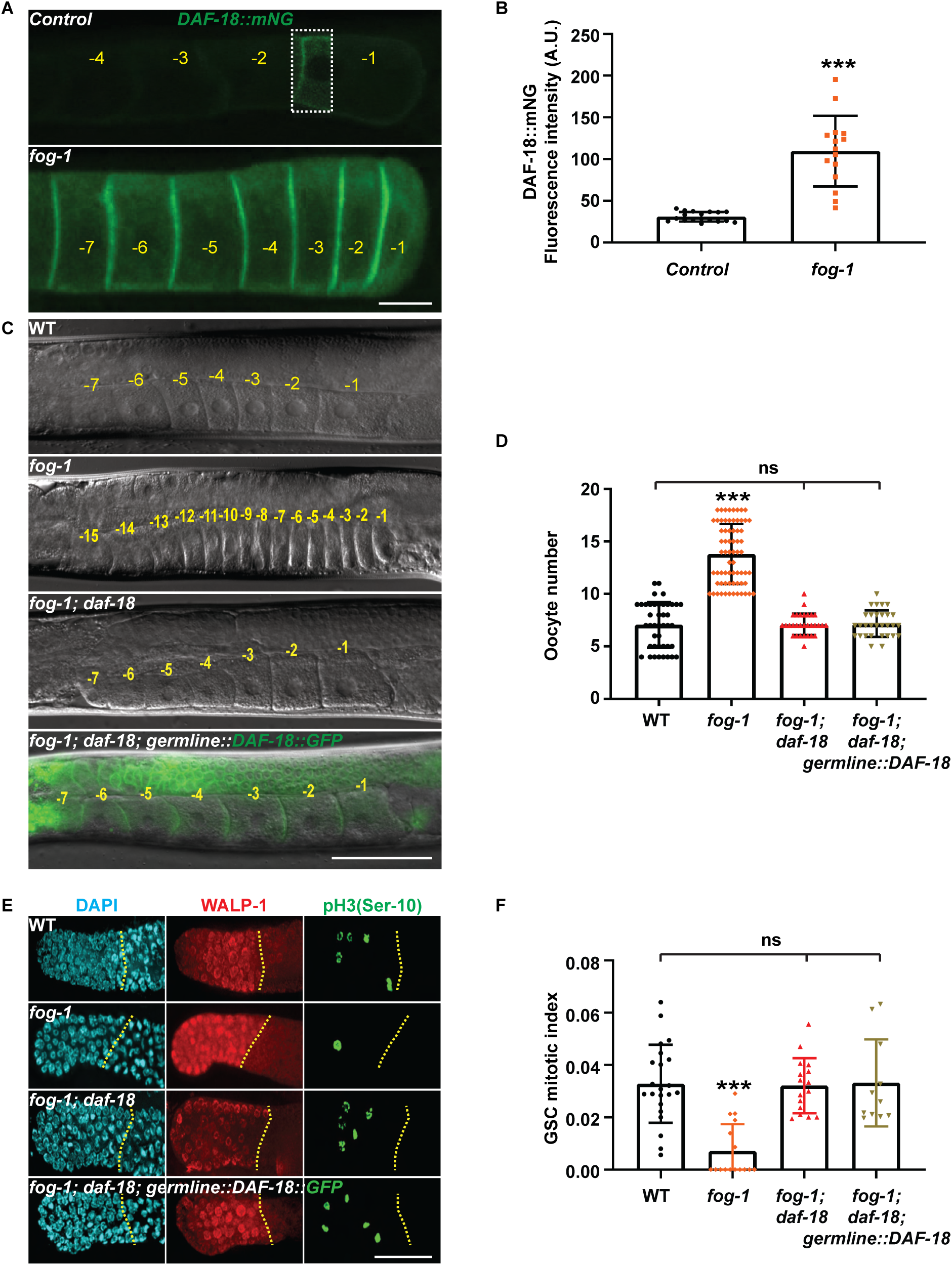
Germline DAF-18 is insufficient to induce oocyte arrest. **(A)** Representative epifluorescence micrographs showing increased levels of endogenous tagged *DAF-18::mNG* at oocyte membranes in *fog-1* day-1 adult hermaphrodites (A1; see methods). Micrographs were acquired using the same parameters, but signal in the boxed region was enhanced in the control for visualization purposes. Negative yellow numbers mark oocytes from distal to proximal. Proximal, right. **(B)** Average *DAF-18::mNG* fluorescence intensity (± standard deviation) at oocyte membranes of A1 animals of the indicated genotypes. Sample sizes: 18, 15. **(C)** Representative differential interference contrast (DIC) or DIC-epifluorescence overlaid micrographs of A1 animals of the indicated genotypes. Negative yellow numbers mark oocytes from distal to proximal. Dorsal, up; proximal, right ventral. **(D)** Average number (± standard deviation) of diakinesis-stage oocytes per gonad arm in A1 hermaphrodites of the indicated genotypes. Sample sizes: 42, 68, 30, 29. **(E)** Representative epifluorescence micrographs of distal germlines dissected from A1 hermaphrodites of the indicated genotypes, stained with DAPI (DNA; blue), anti-WALP-1 (proliferation maker; red) and anti-phospho[ser10] histone H3 (G_2_/M-phase marker; green). Distal, left. Yellow dotted lines, proliferative zone (PZ) boundary. (**A, C, E**) Scale bars: 50 µm. **(F)** Average GSC MIs of A1 hermaphrodites of the indicated genotypes. Sample sizes: 22, 16, 16, 11. **(B, D, F)** Asterisks indicate statistical significance to all other samples. ns, not significant. **(A-F)** Alleles: *fog-1(q253)*, *daf-18(ok480)*, *narSi5[Pmex-5::GFP::DAF-18(+) + unc-119(+)]*.

To validate that the *germline::DAF-18* rescue construct is functional and confirm these surprising results, we asked if it rescued dauer development in *daf-2; daf-18(*ø) Daf-d mutants, since maternally-provided DAF-18 is sufficient to promote dauer entry (Gil et al., 1999). As expected, *germline::DAF-18* maternally rescued dauer formation in a *daf-2; daf-18(*ø) background, in an allelic dosage-dependent manner, while zygotic germline expression was insufficient to promote dauer entry (Figures S1A-S1D). *Germline::DAF-18* did however not prevent the GSC overproliferation that occurs during dauer development in *daf-18(*ø) mutants (Figure S1D), something that requires somatic gonad DAF-18 (Tenen and Greenwald, 2019). These results are consistent with germ-specific transgenic expression and establish that the oocyte quiescence defect of *daf-18(*ø) mutant adults is strictly zygotic. Since a maternal contribution from *germline::DAF-18* to all somatic tissues was insufficient to ensure oocyte arrest (Figures 1C-1F), we conclude that oocyte arrest in the absence of sperm requires significant levels of DAF-18 protein within somatic tissues.

### Muscle DAF-18 is sufficient for oocyte arrest

To identify the somatic tissue(s) from which *daf-18* promotes oocyte arrest in the absence of sperm, we generated new transgenes to restore DAF-18 activity in specific somatic tissues within *fog-1; daf-18(*ø) animals. We used the *unc-54, nhr-72, elt-7,* and *unc-119* promoters to specifically drive *daf-18* expression in non-pharyngeal muscles, seam cells, intestine and nervous system, respectively (Maduro and Pilgrim, 1995; Masse et al., 2005; Miyabayashi et al., 1999; Okkema et al., 1993). Expression of DAF-18 in muscles, henceforth *muscle::DAF-18*, restored oocyte arrest and accumulation, while expression in hypodermal seam, intestinal or neuronal cells had no effect (Figures 2A and 2B). *Muscle::DAF-18* also reinstated GSC quiescence in the absence of sperm (Figures 2C and 2D). These results were confirmed with a second muscle-specific promoter (*Pmyo-3*) (Okkema et al., 1993) (Figures 2A-D). We therefore conclude that, in the absence of sperm, the presence of DAF-18 in non-pharyngeal muscles non-autonomously ensures oocyte arrest and accumulation, along with the concomitant downregulation of GSC proliferation.

**Figure 2.**
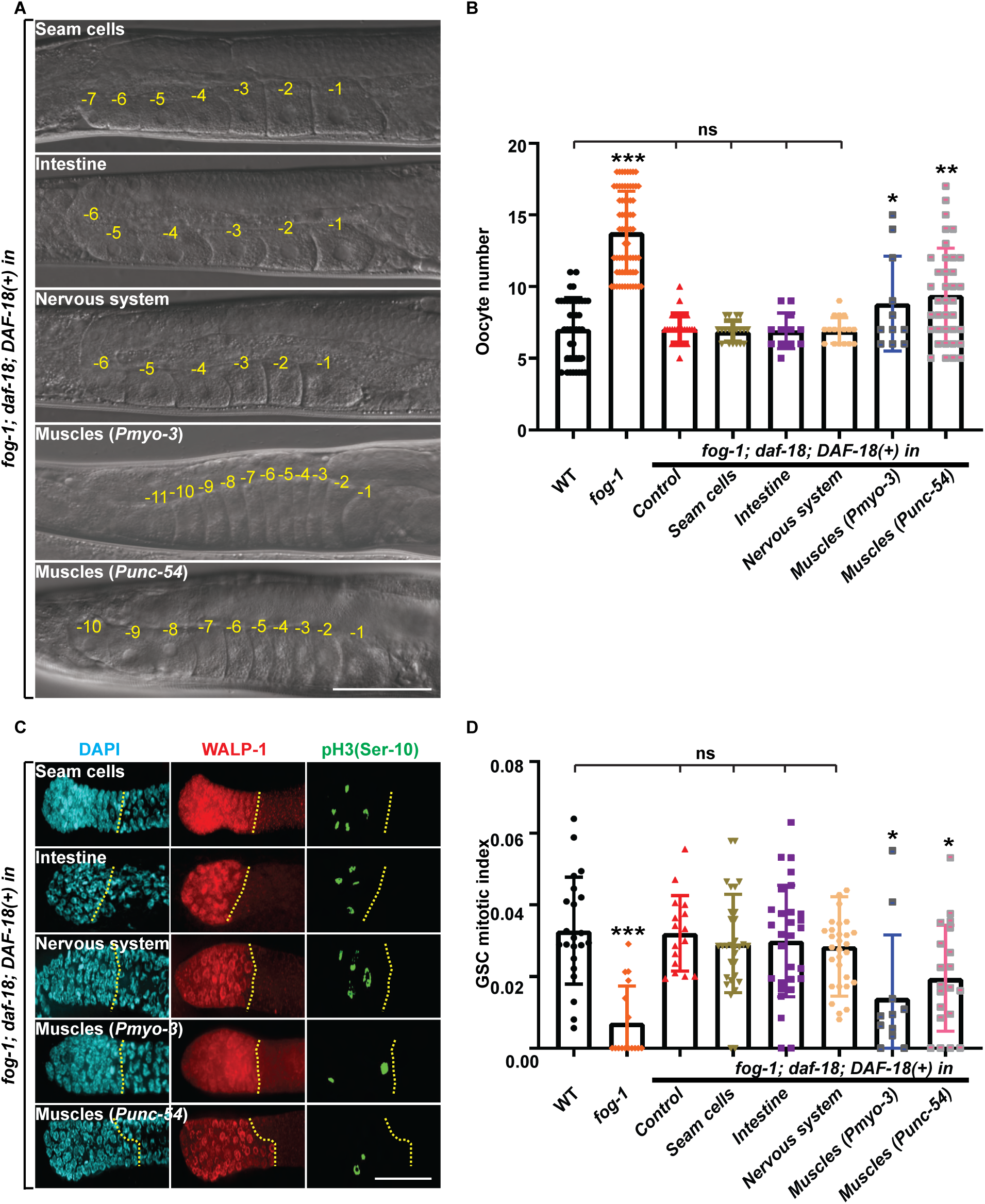
Muscle DAF-18 is sufficient to induce oocyte arrest. **(A)** Representative DIC micrographs of A1 germlines of the indicated genotypes. Negative yellow numbers mark oocytes from distal to proximal. **(B)** Average number (± standard deviation) of diakinesis-stage oocytes per gonad arm in A1 hermaphrodites of the indicated genotypes. Dorsal, up; proximal, right ventral. Sample sizes: 42, 68, 30, 19, 12, 17, 11, 37. **(C)** Representative epifluorescence micrographs of distal gonads isolated from A1 hermaphrodites of the indicated genotypes. Gonads were stained with DAPI (DNA; blue), anti-WALP-1 (proliferation marker; red) and anti-phospho[ser10] histone H3 (G_2_/M-phase marker; green). Distal, left. Yellow dotted lines, PZ boundary. (**A, C**) Tissue-specific promoters: non-pharyngeal muscles (*Pmyo-3*, *Punc-54*), seam cells (*Pnhr-72*), intestine (*Pelt-7*), nervous system (*Punc-119*). Scale bars: 50 µm. **(D)** Average GSC MIs of A1 hermaphrodites of the indicated genotypes. Sample sizes: 22, 16, 16, 31, 29, 30, 11, 20. **(B, D)** Asterisks indicate statistical significance versus *fog-1; daf-18* control. ns, not significant. **(A-D)** Alleles: *fog-1(q253)*, *daf-18(ok480)*, *narEx76[Pnhr-72::DAF-18(+); Pmyo-3::mCherry], narEx64[Pelt-7::DAF-18(+); Pmyo-3::mCherry], narEx85[Punc-119::DAF-18(+); Pmyo-3::mCherry], narEx100[Punc-54::DAF-18(+); Punc-54::GFP], narEx93[Pmyo-3::RFP::DAF-18]*.

### Sp neck DAF-18 is sufficient for oocyte arrest

The *C. elegans* adult hermaphrodite muscular system consists of 20 pharyngeal and 95 body wall muscle cells, in addition to a few other specialized muscles (Altun and Hall, 2009). While both the *myo-3* and *unc-54* promoters are not active in pharyngeal muscles (Ardizzi and Epstein, 1987; Miller et al., 1986), it was conceptually difficult to hypothesize how body wall muscles could promote oocyte arrest in the absence of sperm. In contrast, the Sp and gonadal sheath cells are smooth muscle-like contractile cells that have been heavily implicated in the control of oocyte maturation and ovulation (McCarter et al., 1999; Miller et al., 2001). We closely examined animals bearing *myo-3* and *unc-54* transgenes and detected expression from both promoters in the uterus (Ut), Sp and sheath cells (Figure S2). To determine whether *daf-18* may act within these contractile gonadal tissues, we first restored DAF-18 specifically in the sheath cells, or in the proximal gonad comprising the Sp and Ut, of *fog-1; daf-18(*ø) animals using the *lim-7* and *fos-1a* promoters, respectively (Figure 3A) (Qin and Hubbard, 2015; Voutev et al., 2009). Only *Sp+Ut::DAF-18* expression restored oocyte accumulation and blocked GSC proliferation (Figures 3B-3E). This indicates that the proximal gonad comprises the main site where DAF-18 activity is required to downregulate ovulation and GSC proliferation in the absence of sperm.

**Figure 3.**
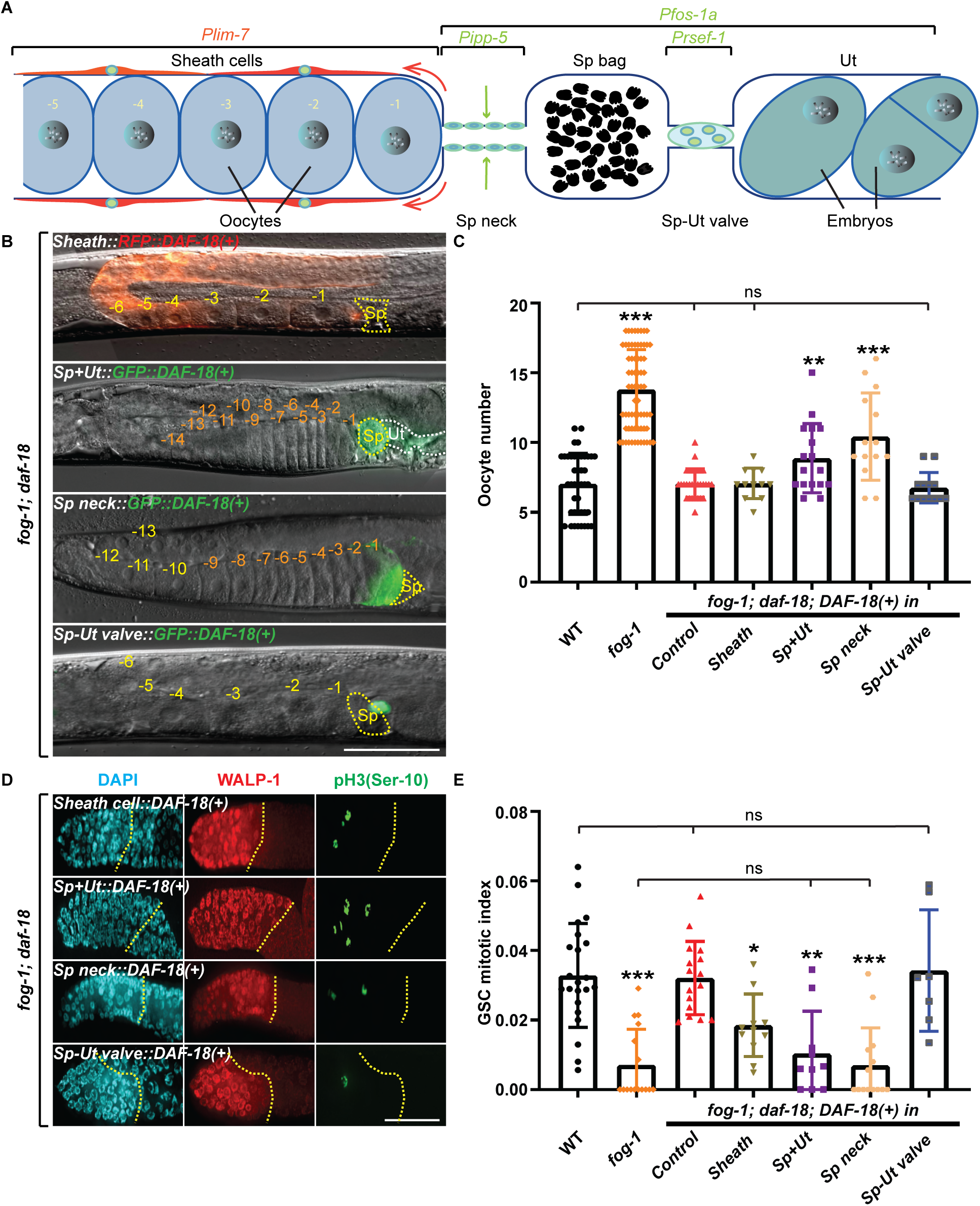
Sp neck DAF-18 is sufficient for oocyte arrest. **(A)** Schematic representation of the *C. elegans* proximal germline and somatic gonad. The Sp consists of three parts: a distal 8-cell neck, central 16-cell bag, and 4-cell syncytial Sp-Ut valve (Castaneda et al., 2020). The sheath cells contract along the distal-proximal axis (red arrows) to pull open the Sp neck around the -1 oocyte. The Sp neck is a sphincter muscle which contraction (blue arrows) resists this opening force (Kelley et al., 2018). Region of expression from tissue-specific promoters are indicated by brackets at the top: sheath cells (*Plim-7*), Sp+Ut (*Pfos-1a*), Sp Neck (*Pipp-5*), Sp-Ut Valve (*Prsef-1*). **(B)** Representative DIC-epifluorescence overlaid micrographs of A1 hermaphrodites of the indicated genotypes. Negative yellow numbers mark oocytes from distal to proximal. Yellow dotted lines, Sp. White dotted line, Ut. Dorsal, up; proximal, right ventral. **(C)** Average number (±standard deviation) of diakinesis-stage oocytes per gonad arm in A1 hermaphrodites of the indicated genotypes. Sample sizes: 42, 68, 30, 12, 13, 17, 14. **(D)** Representative epifluorescence micrographs of distal gonads isolated from A1 hermaphrodites of the indicated genotypes. Gonads were stained with DAPI (DNA; blue), anti-WALP-1 (proliferation maker; red) and anti-phospho[ser10] histone H3 (G_2_/M-phase marker; green). Distal, left. Yellow dotted lines, PZ boundary. **(E)** Average GSC MIs of A1 hermaphrodites of the indicated genotypes. Sample sizes: 22, 16, 16, 12, 7, 10, 14. **(B, D)** Scale bars: 50 µm. **(C, E)** Asterisks indicate statistical significance versus *fog-1; daf-18*. ns, not significant. **(B-E)** Alleles: *fog-1(q253), daf-18(ok480)*, *narEx99[Plim-7::RFP::DAF-18; Pmyo-2::GFP], narEx2[Pfos-1a::GFP::DAF-18; Pmyo-3::mCherry], narEx105[Ptag-312::GFP::daf-18; Pmyo-2::GFP], narEx107[Pipp-5::GFP::daf-18; Pmyo-2::GFP]*.

Interestingly, while *sheath::DAF-18* had no effect on oocyte accumulation, it partially rescued GSC downregulation (Figures 3B-3E). This mild decrease in GSC proliferation in the absence of oocyte accumulation suggests that sheath cells are not the primary site for DAF-18 function. Moreover, as sheath MPK-1 activity can promote GSC proliferation (Robinson-Thiewes et al., 2021), while DAF-18 suppresses MPK-1 activity (Nakdimon et al., 2012; Narbonne et al., 2017; Suzuki and Han, 2006), this partial rescue likely results from the inhibition of sheath MPK-1 by DAF-18 overexpression.

The Sp is the site of fertilization, and consists of three parts: an 8-cell distal neck, a 16-cell central bag, and a syncytial 4-cell Sp-Ut valve (Figure 3A) (McCarter et al., 1997). To materialize ovulation, the gonadal sheath cells enter in a tug-of-war with the Sp neck until they successfully pull it open around the proximal (-1) oocyte (Castaneda et al., 2020). Once the oocyte enters the Sp, it is immediately fertilized, and the Sp neck and bag then contract to push the egg through the Sp-Ut valve and into the Ut (Video S1) (Castaneda et al., 2020; Yamamoto et al., 2006). Two sphincter muscles, the Sp neck and the Sp-Ut valve, can therefore block the passage of oocytes and promote their accumulation in the absence of sperm. We therefore specifically restored DAF-18 either in the Sp neck, or the Sp-Ut valve, of *fog-1; daf-18(*ø) animals, using the *ipp-5* and *rsef-1* promoters, respectively (Bui and Sternberg, 2002; Ghosh and Sternberg, 2014). *Sp neck::DAF-18* rescued both oocyte arrest and GSC proliferation, while *Sp-Ut valve::DAF-18* did not (Figures 3B-3E). We conclude that DAF-18 expression in the Sp neck, formed by 8 myoepithelial cells, is sufficient to non-autonomously promote oocyte arrest and accumulation in the absence of sperm, and to induce the concomitant downregulation in GSC proliferation.

### DAF-18’s lipid phosphatase activity is required for oocyte arrest

DAF-18 is a dual-specificity phosphatase that could block ovulation through its lipid or protein phosphatase activity, or through a catalytically independent function. To identify the key function, we introduced a G174E (equivalent to human G129E) substitution in endogenous DAF-18 to specifically remove its lipid phosphatase activity (Myers et al., 1997; Nakdimon et al., 2012). We found that *fog-1; daf-18(G174E)* mutants wasted their oocytes like *fog-1; daf-18(*ø) (Figures 4A and 4B) while their GSCs showed sustained proliferation (Figures 4C and 4D). Without ruling out the possibility of an additional protein phosphatase requirement, these results show that the catalytic function of DAF-18, including its lipid phosphatase activity, is required to prevent ovulation in the absence of sperm.

**Figure 4.**
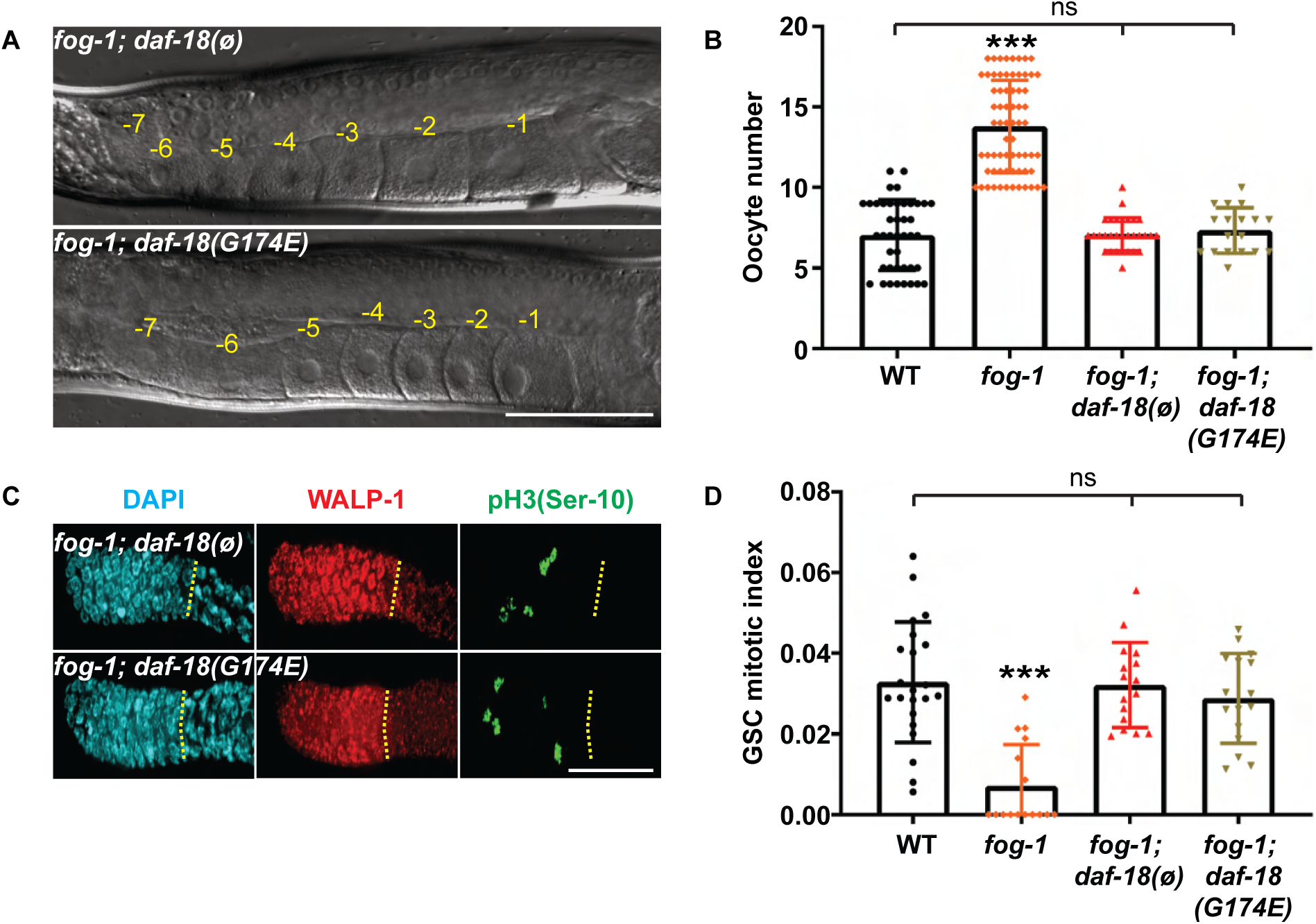
DAF-18’s lipid phosphatase activity induces oocyte arrest. **(A)** Representative DIC micrographs of A1 germlines of the indicated genotypes. Negative yellow numbers mark oocytes from distal to proximal. Dorsal, up; proximal, right ventral. **(B)** Average number (± standard deviation) of diakinesis-stage oocytes per gonad arm in A1 hermaphrodites of the indicated genotypes. Sample sizes: 42, 69, 30, 18. **(C)** Representative distal germlines dissected from A1 animals of the indicated genotypes, stained with DAPI (DNA; blue), anti-WALP-1 (proliferation maker; red) and anti-phospho[ser10] histone H3 (G_2_/M-phase marker; green). Distal, left. Yellow dotted lines, PZ boundary. (**A, C**) Scale bars: 50 µm. **(D)** Average GSC MIs of A1 hermaphrodites of the indicated genotypes. Sample sizes: 22, 16, 16, 16. **(B, D)** Asterisks indicate statistical significance *versus* all other samples. ns, not significant. **(A-D)** Alleles: *fog-1(q253), daf-18(ok480), daf-18(nar57[G174E])*.

### DAF-18 prevents unwanted ovulation independently of PIP_2_ and PLC-1/ITR-1/Ca^2+^ signaling

The Sp neck is a sphincter muscle that is constricted between ovulation events to block the premature entrance of the maturing proximal oocyte, despite ongoing basal sheath cell contractions (Aono et al., 2004; Castaneda et al., 2020; Govindan et al., 2009; Yamamoto et al., 2006). The final maturation of the proximal oocyte is however thought to provoke the release of the epidermal growth factor (EGF) LIN-3 and trigger stronger ovulatory sheath contractions, which eventually overcome those at the Sp neck and force it open as they pull it around the large oocyte that is now fertilization-competent (Clandinin et al., 1998; Yin et al., 2004). Contractility in both sheath and Sp is dependent on actin/myosin interactions, which are stimulated by rises in cytoplasmic Ca^2+^ concentrations following its release from the endoplasmic reticulum by the inositol triphosphate (IP_3_) receptor ITR-1 (Castaneda et al., 2020; Clandinin et al., 1998; Kovacevic et al., 2013). DAF-18’s main lipid phosphatase activity consists in dephosphorylating PIP_3_ into PIP_2_, which is a substrate for the phospholipase C-ε PLC-1. PLC-1 cleaves PIP_2_ to generate inositol 1,4,5-triphosphate (IP_3_), which in turn favors the opening of the IP_3_-sensing ITR-1 Ca^2+^ channel to release Ca^2+^ in the cytoplasm and promote contractility (Kovacevic et al., 2013). To test whether DAF-18 promotes Sp neck contractility through regulation of PIP_2_ levels and PLC-1/ITR-1/Ca^2+^ signaling, we first recorded time-lapse sequences of ovulating animals expressing the Ca^2+^ fluorescent sensor GCaMP3 in the Sp (Figure 5A) (Bouffard et al., 2019). This genetically encoded cytoplasmic Ca^2+^ sensor is a synthetic fusion between GFP, calmodulin and the M13 peptide that fluoresces green only when Ca^2+^ bound (Figure 5B) (Nakai et al., 2001). In control hermaphrodites, as expected, the Sp neck was contracted before ovulation, relaxed as it was stretched-open and pulled around the mature oocyte by the sheath cells during ovulation, and then contracted again, this time stronger and together with central bag cells, to push the fertilized oocyte through the Sp-Ut valve (Figure 5A; Video S1). In spermless *fog-1* mutants however, the Sp neck always remained constricted, and effectively prevented oocytes to enter the Sp (Figure 5A; Video S2). In *fog-1; daf-18(*ø) animals, the Sp neck relaxed normally to allow oocyte entry, after which the neck and bag contracted to expulse it into the Ut, as in control animals. (Figure 5A; Video S3). Interestingly, oocyte Sp entry time was significantly shorter in *fog-1; daf-18(*ø) mutants than in controls, consistent with a lower Sp neck dilation resistance (Figure S3A). We conclude that DAF-18 cell autonomously prevents Sp neck dilation in the absence of sperm signals (Miller et al., 2001).

**Figure 5.**
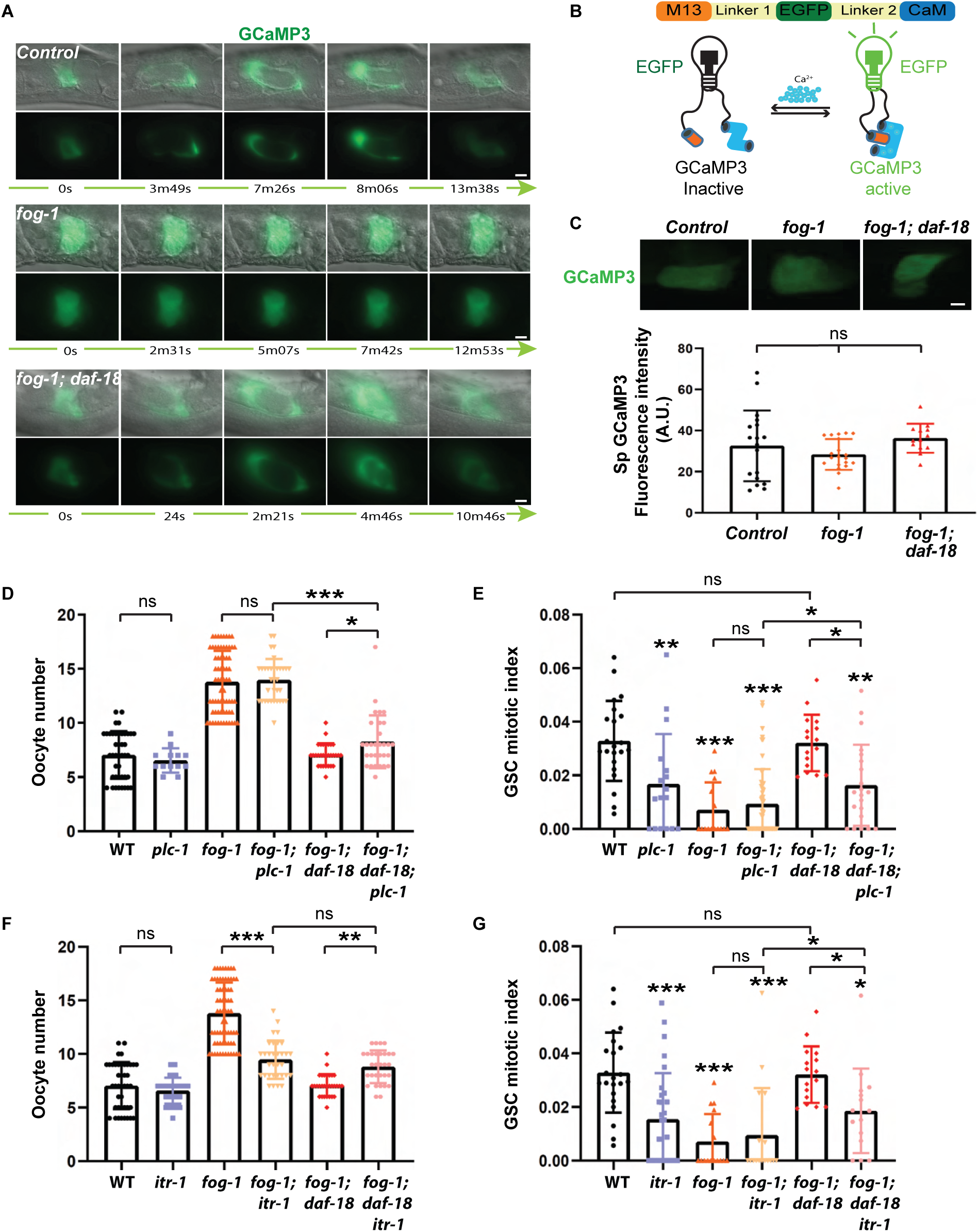
DAF-18 prevents unwanted ovulation independently of PLC-1/ITR-1/Ca^2+^ signaling. **(A)** Representative time frames showing Sp Ca^2+^ flows during key ovulatory events in A1 hermaphrodites of the indicated genotypes. Videos are available online (Videos S1-S3). Sample sizes: 17, 50, 9. **(B)** Schematic representation of the GCaMP3 Ca^2+^ sensor, a synthetic fusion between EGFP, calmodulin (CaM) and the M13 peptide. When Ca^2+^ bound, the CaM domain undergoes a conformational change that enables its interaction with M13’s alpha helix, resulting in bright fluorescence of the GCaMP3 sensor (Iseppon et al., 2022). **(C)** Quantitative GCaMP3 fluorescence analysis showed that average (± standard deviation) Sp cytoplasmic Ca^2+^ concentrations were unaffected in *fog-1* and *fog-1; daf-18(*ø) mutants. Sample sizes: 18, 20, 13. (**A, C**) Distal, left. Scale bars: 50 µm. **(D)** Average number (± standard deviation) of diakinesis-stage oocytes per gonad arm in A1 hermaphrodites of the indicated genotypes. Sample sizes: 42, 13, 68, 34, 30, 18. **(E)** Average GSC MIs (± standard deviation) of A1 hermaphrodites of the indicated genotypes. Sample sizes: 22, 17, 16, 53, 16, 21. **(F)** Average number (± standard deviation) of diakinesis-stage oocytes per gonad arm in A1 hermaphrodites of the indicated genotypes. Sample sizes: 42, 40, 68, 32, 30, 30. **(G)** Average GSC MIs (± standard deviation) of A1 hermaphrodites of the indicated genotypes. Sample sizes: 22, 30, 16, 18, 16, 15. **(C-G)** Asterisks indicate statistical significance versus wild-type or as indicated by brackets. ns, not significant. **(A-G)** Alleles*: fog-1(q253), daf-18(ok480), itr-1(sa73)*, *xbIs1101[Pfln-1::GCaMP], plc-1(rx1)*, *kfEx2[plc-1(+); Psur-5::GFP]*.

We next asked whether DAF-18 increases Sp neck contractility by rising cytoplasmic Ca^2+^ concentrations. We therefore measured inter-ovulatory Sp GCaMP3 average fluorescence intensities as a proxy for Sp intracellular Ca^2+^ concentration. Sp GCaMP3 fluorescence intensities were however undistinguishable in control, *fog-1* and *fog-1; daf-18(*ø) animals (Figure 5C). These results suggest that DAF-18 promotes Sp neck contractility independently of Sp Ca^2+^ flows.

Since the loss of *daf-18* is expected to reduce PIP_2_ levels, which in turn forecasts reduced PLC-1 outputs (Kariya et al., 2004), we still investigated the interaction between *daf-18* and *plc-1*. In the presence of sperm, the loss of *plc-1* causes the trapping of embryos in the Sp, since its neck and bag lack the ability to strongly contract and expulse eggs through the Sp-Ut valve (Castaneda et al., 2020), which does not express *plc-1* (Kariya et al., 2004). As expected, the *fog-1; plc-1(ø)* double mutant exhibited lower contractility in its Sp neck and bag, resulting in oocyte trapping (Figure S3B), reminiscent of the *plc-1(ø)* phenotype (Kariya et al., 2004). If DAF-18 were to affect Sp neck contractility exclusively through PLC-1, its removal should not exacerbate the *plc-1(ø)* phenotype. However, we found that the *fog-1; daf-18(ø); plc-1(ø)* triple mutants had clearly more oocytes trapped within their Sp than *fog-1; plc-1(*ø) doubles (Figures S3B and S3C). We also quantified Sp GCaMP3 fluorescence intensity in *fog-1; plc-1(ø*) doubles and *fog-1; daf-18(ø); plc-1(ø)* triples and did not detect any difference (Figure S3C), confirming that the loss of DAF-18 does not affect Sp Ca^2+^ levels, even in the absence of PLC-1. Most strikingly, the loss of *daf-18* still largely prevented oocyte accumulation in the absence of *plc-1* (Figure 5D), suggesting that it fulfills this role largely independently from *plc-1*. We attribute the weak rescue of anovulated oocyte accumulation to the introduction of a new blockade in the path of oocytes in the absence of *plc-1*, this time at the Sp-Ut valve, that may have caused oocytes to backlog into the proximal gonad once the Sp got maximally stretched (Figures 5D and S3B). Regarding GSC proliferation, the loss of *plc-1* had a negative effect on the GSC MI on its own, but it did not add to the negative homeostatic pressure present in *fog-1* mutants (Figure 5E). Germline feminization and the loss of *plc-1* could therefore negatively regulate GSC proliferation through the same downstream mechanism. Interestingly, *fog-1; daf-18(ø); plc-1(*ø) triple mutants had a GSC MI that was intermediate between the *fog-1;plc-1(*ø) and the *fog-1; daf-18(ø)* doubles (Figure 5E), indicating that *daf-18* suppresses GSC proliferation in feminized mutants independently from *plc-1*.

To further explore the interaction between *daf-18* and Ca^2+^ signaling, we reduced the activity of the *C. elegans* unique IP_3_ receptor that releases Ca^2+^ into the cytoplasm downstream of PLC proteins, ITR-1 (Baylis et al., 1999), using the viable *itr-1(sa73)* reduction-of-function *(rf)* allele (Dal Santo et al., 1999). We first note that feminized *itr-1(rf)* mutants had lower Sp intracellular Ca^2+^ levels than feminized *plc-1(*ø), but that this was again, not modified by the additional loss of *daf-18* (Figure S3B). As *plc-3* is expressed in the spermatheca in addition to *plc-1* (Hiatt et al., 2009; Yin et al., 2004), we attribute this result to the loss of all PLC-dependent Ca^2+^ influx when *itr-1* is mutated. Otherwise, this *itr-1(rf)* allele interacted with *daf-18(ø)* in a manner similar to *plc-1(ø)* in all assays (Figures 5F, 5G, S3B and S3C) except for one notable difference. Namely, and potentially due to their presumably looser Sp neck (Figure S3B), *fog-1; itr-1(rf)* doubles did not accumulate as many anovulated oocytes as *fog-1* singles (Figures 5F and S3B). As a result, the removal of *daf-18* in the *fog-1; itr-1(rf)* background had no additional effects on oocyte accumulation. But as seen with *plc-1(ø)*, *daf-18* promoted GSC quiescence independently from *itr-1* (Figure 5G). Altogether, these data indicate that DAF-18 prevents unwanted ovulation and promote GSC quiescence in the absence of sperm largely independently from PLC-1/ITR-1/Ca^2+^ signaling.

### DAF-18 prevents ovulation by reducing Sp PIP_3_ levels and AKT-1/2 activity

If Ca^2+^ signaling is largely unmoved downstream of DAF-18 lipid phosphatase activity’s main product, PIP_2_, we reasoned that the ovulation defects may instead result from changes in DAF-18’s substrate levels, PIP_3_. We therefore used anti-PIP_3_ antibodies to evaluate their levels in the Sp. In wild-type gonads, there was relatively little PIP_3_ throughout the germline, distal tip and sheath cells, while quite strikingly, much higher PIP_3_ levels were associated with Sp cell membranes (Figure 6A). Feminization did not perturb Sp PIP_3_ levels but there were more PIP_3_ in *daf-18(*ø) Sp, both in control and feminized hermaphrodites (Figures 6B and 6C). Therefore, DAF-18 activity lowers Sp PIP_3_ levels, which may in turn suppress AKT-1/2 (the *C. elegans* AKT/PKB orthologs) through reducing PDK-1 activation.

**Figure 6.**
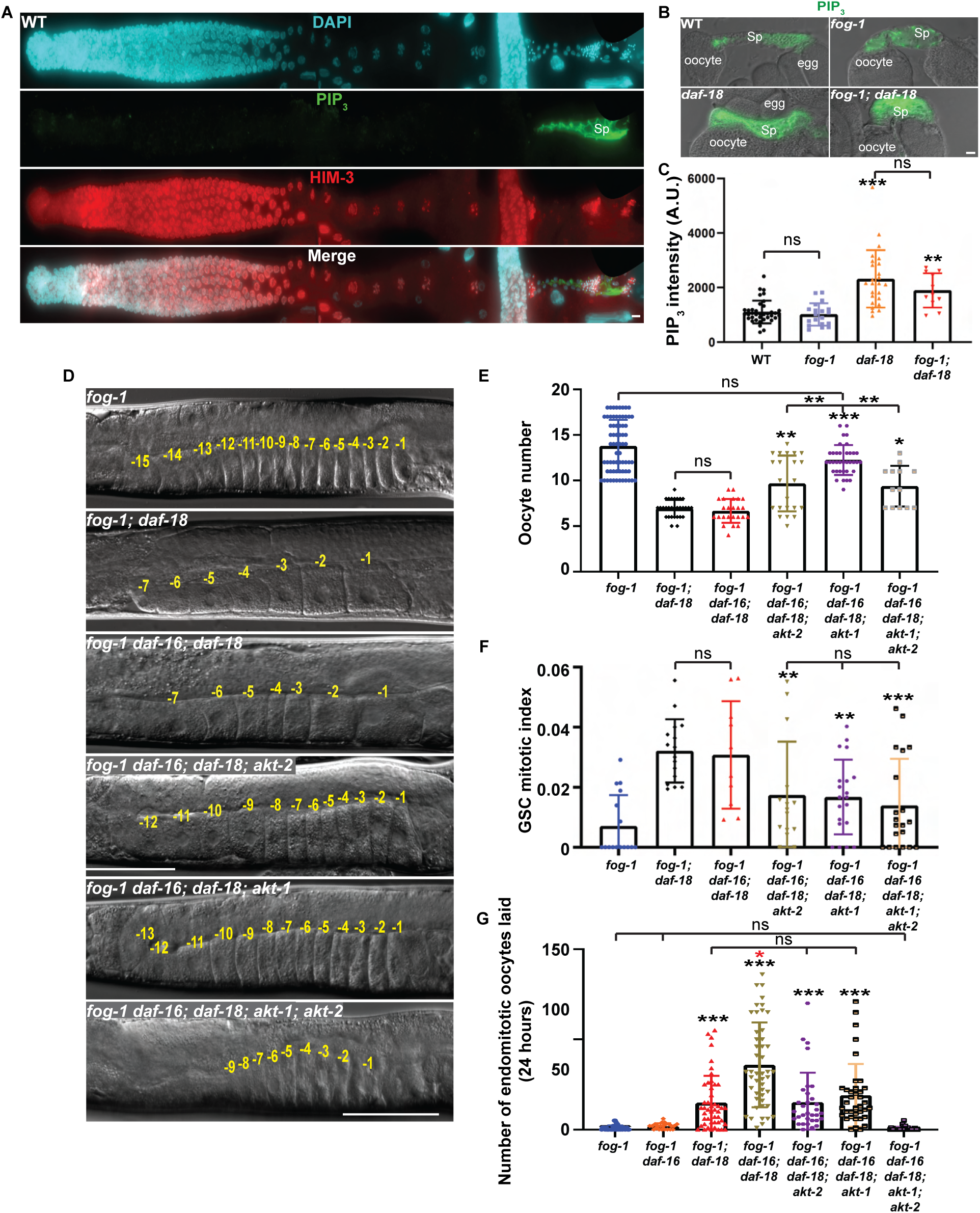
DAF-18 prevents unwanted ovulation by inhibiting AKT-1/2. **(A)** Representative epifluorescence micrographs of a whole germline dissected from a wild-type A1 hermaphrodite stained with DAPI (blue), anti-PIP_3_ (green), and anti-HIM-3 (differentiation marker, red). PIP_3_ is dramatically enriched at the membranes of Sp cells. Distal, left. **(B)** Representative DIC images of fixed and dissected proximal germlines from A1 animals of the indicated genotypes, stained with anti-PIP_3_ (green) and overlaid with the resulting epifluorescence signals. **(C)** Average Sp PIP_3_ levels in hermaphrodites of the indicated genotypes. Sample sizes: 38, 25, 19, 11. **(D)** Representative DIC micrographs of A1 germlines of the indicated genotypes. The DIC micrographs of *fog-1* and *fog-1; daf-18(*ø) controls are duplicates of Figure 1C. Negative yellow numbers mark oocytes from distal to proximal. Dorsal, up; proximal, right ventral. (**A, B, D**) Scale bars: 50 µm. **(E)** Average number (± standard deviation) of diakinesis-stage oocytes per gonad arm in A1 hermaphrodites of the indicated genotypes. Sample sizes: 68, 30, 24, 22, 36, 13. **(F)** Average GSC MIs (± standard deviation) of A1 hermaphrodites of the indicated genotypes. Sample sizes: 16, 16, 10, 15, 20, 20. **(G)** Average number (± standard deviation) of endomitotic oocytes laid by animals of the indicated genotypes over a period of 24 hours after A1. Sample sizes: 40, 22, 47, 47, 32, 37, 34. **(C,E-G)** Asterisks indicate statistical significance versus (**C**) wild-type for *daf-18* and *fog-1* for *fog-1; daf-18*, (**E-F**) *fog-1; daf-18(*ø) controls and (**G**) *fog-1* controls, or as indicated by brackets. The red asterisk represents statistical significance versus all other groups. ns, not significant. **(A-G)** Alleles*: fog-1(q253), daf-16(mu86), daf-18(ok480), akt-1(ok525), akt-2(ok393)*.

Incidentally, *akt-1* and *akt-2* are highly expressed in the Sp (Padmanabhan et al., 2009), suggesting that they may play a role in this tissue. However, neither *akt-1* nor *akt-2* single null mutations significantly affected oocyte numbers or the GSC MI (Figures S4A-S4C). To investigate a potential redundancy, *akt-1/2(ø)* doubles were combined with *daf-16(ø)* to prevent constitutive dauer formation and allow triples to grow into adults (Ogg et al., 1997). Both *fog-1 daf-16(ø)* doubles and *fog-1 daf-16(ø); akt-1/2(ø)* quadruples had oocyte numbers and GSC proliferation rates that matched those of *fog-1* singles (Figures S4A-C). We conclude that *akt-1/2* do not significantly influence ovulation and GSC proliferation in feminized hermaphrodites when *daf-18* is present and *daf-16* is absent.

Next, we investigated the loss of *akt-1/2* in the absence of *daf-18*. The removal of *daf-16* from *fog-1; daf-18(ø)* did not perturb proximal oocyte numbers and the GSC MI (Figures 6D-F). We removed *akt-1* and/or *akt-2* from this strain to generate *fog-1 daf-16(ø); daf-18(ø); akt-1(ø)* and *fog-1 daf-16(ø); daf-18(ø); akt-2(ø)* quadruples, as well as the *fog-1 daf-16(ø); daf-18(ø); akt-1/2(ø)* quintuple. Both quadruple mutants had significantly more oocytes and a lower GSC MI than *fog-1; daf-18(ø),* and *fog-1 daf-16(ø); daf-18(ø)* controls (Figures 6D-F), indicating that the loss of either *akt-1* or *akt-2* significantly rescues both *daf-18(ø)* defects. The rescuing effect of *akt-1(ø)* on oocyte accumulation was significantly greater than that of *akt-2(ø)* (Figure 6E), suggesting a more prominent role for AKT-1 in suppressing unwanted ovulation. Consistent with this, Sp AKT-1::GFP levels are visibly higher than those of AKT-2::GFP (Padmanabhan et al., 2009). Although the number of anovulated oocytes present in the proximal gonad of *fog-1 daf-16(ø); daf-18(ø); akt-1(ø)* quadruple mutant was undistinguishable from *fog-1* controls (Figures 6D and 6E), the quadruple mutant laid more endomitotic oocytes than *fog-1* controls (Figure 6G), indicating that some level of unwanted ovulation still occurred.

Because of the partial suppression observed in each quadruple mutant, we expected the simultaneous removal of *akt-1* and *akt-2* to fully restore oocyte accumulation and GSC downregulation in feminized *daf-18(ø)* hermaphrodites. However, we found that the *fog-1 daf-16(ø); daf-18(ø); akt-1/2(ø)* quintuple mutant accumulated more oocytes than *fog-1; daf-18(ø)* and *fog-1 daf-16(ø); daf-18(ø)* controls, but still not as many as *fog-1* singles (Figures 6D and 6E). In fact, the rescue of oocyte accumulation was better in the *fog-1 daf-16(ø); daf-18(ø); akt-1(ø)* quadruple than in the *fog-1 daf-16(ø); daf-18(ø); akt-1/2(ø)* quintuple (Figures 6D and 6E). We nonetheless note that, for unknown reasons, the *fog-1 daf-16(ø); daf-18(ø); akt-1/2(ø)* quintuple mutant appeared somewhat sickly and grew slowly (Figure S4D), which may explain the observed partial suppression. Importantly however, the quintuple mutant completely stopped to futilely lay endomitotic oocytes (Figure 6G). These results suggest that *akt-1* and *akt-2* work additively to permit ovulation in the absence of sperm and *daf-18(*ø), and that they are together sufficient to mediate these unwanted ovulation events. We therefore conclude that the loss of DAF-18 elevates PIP_3_ levels in the Sp neck, where they activate AKT-1/2 which, in turn and independently from DAF-16, promote Sp neck dilation to allow ovulation in the absence of sperm.

### AKT-1/2 likely phosphorylate RHO-1 to facilitate Sp neck dilation and ovulation

Sp contractions are triggered by actin-myosin interactions, the efficiency of which is increased by rises in intracellular Ca^2+^ concentrations. Namely, intracellular Ca^2+^ activates a Ca^2+^-dependent myosin light chain (MLC) kinase that phosphorylates MLC to promote its interaction with actin (Kelley et al., 2018). Contractility is further modulated by Rho kinase (ROCK) signaling, in which the immediate upstream activator RHO-1/RhoA is activated by guanine nucleotide exchange factors (GEFs) and inhibited by GTPase-activating proteins (GAPs) (Castaneda et al., 2020; Kalpachidou et al., 2019; Kariya et al., 2004; Kovacevic et al., 2013; Lundquist, 2006; Tan and Zaidel-Bar, 2015). As such, GTP-bound active RHO-1 interacts with LET-502/ROCK, which directly phosphorylates both MLC and MLC phosphatase to increase Ca^2+^ sensitivity (Kovacevic et al., 2013; Tan and Zaidel-Bar, 2015). As the loss of *daf-18* does not perturb Sp Ca^2+^ levels (Figures 5 and S3), we reasoned that AKT-1/2 could affect contractility through ROCK without perturbing Ca^2+^ signaling. We further deduced that AKT-1/2 were likely to act at the level of RHO-1, or downstream, since upstream components (KIN-1 and GSA-1) also influence Ca^2+^ signaling (Castaneda et al., 2020).

Interestingly, we identified a single perfectly conserved AKT RXRXXS consensus phosphorylation motif (Paradis and Ruvkun, 1998) around RHO-1’s serine 73 (S73), also S73 in human RhoA (Figure 7A), but no such motif in LET-502. We utilized the High Ambiguity Driven protein-protein DOCKing (HADDOCK) 2.4 (de Vries et al., 2010) platform to model a potential physical interaction between the AKT-1 kinase domain and RHO-1’s S73 and obtained a docking score predictive of its likelihood (Figures 7B-D; Video S4). To have a useful comparison, we repeated this *in silico* experiment between AKT-1 and DAF-16 to obtain the docking scores for each of its four well-established AKT target sites (Paradis and Ruvkun, 1998). The docking score for AKT-1 and RHO-1’s S73 was highly negative indicating that the interaction is favorable, with a value comprised within the range of those obtained for the DAF-16 sites (Figure 7B). Template-free docking scores between AKT-1 and RHO-1 or DAF-16, using the HDOCK platform (Yan et al., 2020), were also negative and similar (Figure S5A). RHO-1’s S73 is therefore a potential direct phosphorylation target of AKT kinases.

**Figure 7.**
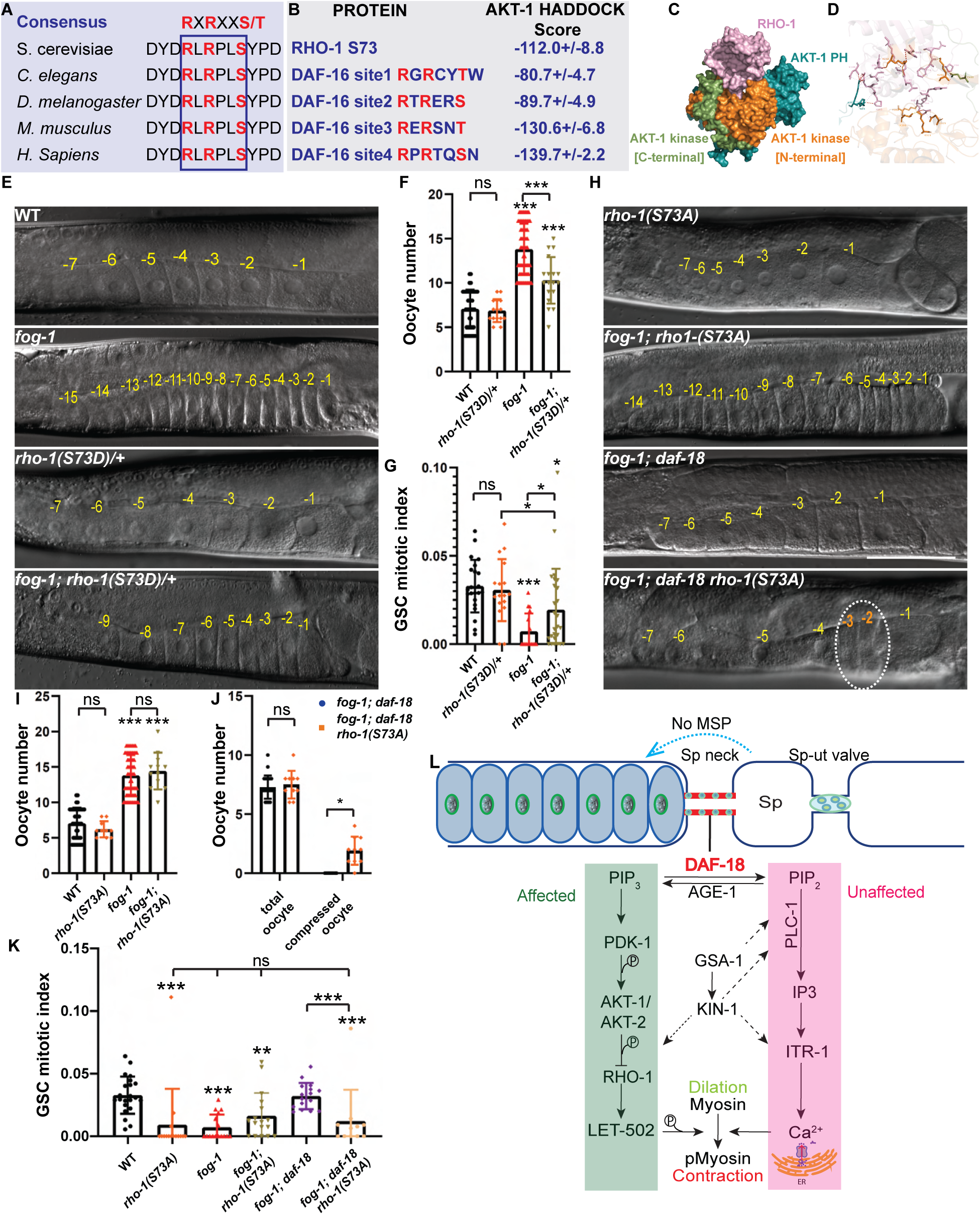
Modifications of RHO-1’s consensus AKT phosphorylation site influence oocyte arrest. **(A)** A single perfectly conserved AKT RXRXXS consensus phosphorylation site was identified within RHO-1. **(B)** The HADDOCK platform was used to predict docking energies between AKT-1’s kinase domain interface and RHO-1(S73), and to compare them with those obtained for AKT-1 and its established target phosphorylation sites on DAF-16 (Paradis and Ruvkun, 1998). **(C)** HADDOCK (v. 2.4) generated AKT-1 (blue: PH domain; orange: kinase N-terminal; green: kinase C-terminal) bound RHO-1 (pink) complex. **(D)** Close-up of the interaction interface showing hydrogen bonds predicted to form between AKT-1 and RHO-1. (**E**) Representative DIC micrographs of A1 germlines of the indicated genotypes. **(F)** Average number (±standard deviation) of diakinesis-stage oocytes per gonad arm in A1 hermaphrodites of the indicated genotypes. Sample sizes: 42, 15, 68, 18. **(G)** Average GSC MIs (±standard deviation) of A1 hermaphrodites of the indicated genotypes. Sample sizes: 22, 17, 16, 26. **(H)** Representative DIC micrographs of A1germlines of the indicated genotypes. White dotted circle, visibly compacted oocytes. **(E, H)** The DIC micrographs of WT, *fog-1* and *fog-1; daf-18(*ø) are duplicates of Figure 1C. Negative yellow numbers mark oocytes from distal to proximal. Dorsal, up; proximal, right ventral. Scale bar: 50 µm. **(I)** Average number (±standard deviation) of diakinesis-stage oocytes per gonad arm in A1 hermaphrodites of the indicated genotypes. Sample sizes: 42, 10, 68, 12. **(J)** Even though the *fog-1; daf-18(*ø*) rho-1(S73A)* strain does not exhibit significant oocyte accumulation, most animals have some visibly compacted oocytes, something that never occurs in *fog-1; daf-18(*ø) doubles. Sample sizes, 25, 10. **(K)** Average GSC MIs (±standard deviation) of A1 hermaphrodites of the indicated genotypes. Sample sizes: 22, 15, 16, 17, 16, 11. **(L)** Model for the non-autonomous regulation of oocyte ovulation by Sp neck DAF-18’s lipid phosphatase activity. Arrows represent activation and bars represent inhibition. Even though the removal of DAF-18 should perturb both substrate (PIP_3_) and product levels (PIP_2_), our data demonstrate that DAF-18 mainly affects ovulation through perturbations in signaling downstream of substrate levels. Namely that the resulting increase in PIP_3_ levels activate AKT-1,2, which can then phosphorylate and inhibit RHO-1 in the Sp neck to suppress contractility and facilitate ovulation. **(F-G, I-K)** Asterisks indicate statistical significance versus wild-type or as indicated by brackets. ns, not significant. **(E-K)** Alleles*: fog-1(q253), daf-18(ok480), rho-1(nar63[S73A]), rho-1(nar64[S73D])/nT1*.

To evaluate whether AKT-1/2 may directly phosphorylate RHO-1 to prevent Sp neck dilation and unwanted ovulation in the absence of sperm, we mutated endogenous RHO-1 S73 to either alanine (A) to prevent phosphorylation, or to aspartic acid (D) to mimic phosphorylation (Bauer et al., 2003; Leger et al., 1997; Peng et al., 2012). Based on the phenotype of *daf-18(ø)* and its suppression by *akt-1/2(ø)*, we predicted that a phospho-defective RHO-1(S73A) would no longer be inhibited by AKT phosphorylation and thus exhibit persistent activity, while a phospho-mimetic RHO-1(S73D) would have reduced activity. Consistent with this, phosphorylation usually has a negative effect on the activity of Rho GTPases (Haga and Ridley, 2016).

Unfortunately, we could not obtain homozygous *rho-1(S73D)* animals as this substitution caused a fully penetrant embryonic lethality (Figure S5B and S5C), similar to the *rho-1(ø)* phenotype (McMullan and Nurrish, 2011; Wernike et al., 2016). This result nonetheless suggests that constitutive phosphorylation at S73 severely impairs RHO-1 activity. We thus examined *fog-1; rho-1(S73D)/+* heterozygotes as their presumably reduced functional RHO-1 levels could promote Sp neck dilation and prevent oocyte accumulation. As a matter of fact, *fog-1; rho-1(S73D)/+* heterozygotes accumulated fewer oocytes than *fog-1* controls (Figure 7E and 7F). Consistent with this, *fog-1; rho-1(S73D)/*+ heterozygotes also had a partially restored GSC MI (Figure 7G). AKT-1/2 phosphorylation on RHO-1(S73) may therefore reduce activity and Sp neck contractility, preventing robust oocyte accumulation and suppression of GSC proliferation in the absence of sperm.

Although they lay much fewer and less viable eggs (Figures S5D and S5E), homozygous *rho-1(S73A)* mutants are viable (Figure 7H). We expected RHO-1(S73A) to prevent AKT-1/2 inhibitory phosphorylation and thus present a persistent activity, but not necessarily increased or constitutive activity. As such, the *rho-1(S73A)* mutation did not perturb oocyte numbers on its own in the presence or absence of sperm (Figures 7H and 7I). We expected *rho-1(S73A)* to suppress the loss of *daf-18* by preventing RHO-1 downregulation by the resulting AKT-1/2 upregulation. However, *fog-1; daf-18(ø) rho-1(S73A)* triple mutants did not accumulate more oocytes than *fog-1; daf-18(ø)* controls (Figures 7H and 7J). They nonetheless often exhibited some visibly compressed oocytes (Figures 7H and 7J), indicating that the Sp neck did oppose an increased resistance. This partial rescue may be linked to the substantial sterility of *rho-1(S73A)* mutants (Figures S5D and S5E), which could be unable to generate oocytes rapidly enough to sustain their accumulation.

Consistent with this, *rho-1(S73A)* had a negative effect on GSC proliferation, suggesting that persistent RHO-1 activity limits GSC proliferation (Figure 7K). Interestingly, feminization did not further reduce GSC proliferation in *rho-1(S73A)* mutants (Figure 7K), raising the possibility that feminization and *rho-1(S73A)* may negatively regulate GSC proliferation through the same downstream mechanism. Moreover, GSC proliferation remained low when *daf-18* was removed from *fog-1; rho-1(S73A)* doubles, indicating that phosphorylation on RHO-1(S73) is required for AKT-1/2 to promote GSC proliferation in *fog-1; daf-18(ø)* mutants (Figure 7K). Altogether, these data support a model in which, in the absence of sperm, DAF-18 is required to prevent AKT-1/2 from depositing an inhibitory phosphate group on RHO-1’s S73, thereby keeping it active to tighten the Sp neck and prevent spontaneous ovulation (Figure 7L).

### DAF-18 acts cell non-autonomously to prevent benign tumorigenesis

Beyond preventing oocyte wastage in the absence of sperm, this newly identified cascade that permits homeostatic control of GSC proliferation may also serve to avert the formation of benign differentiated tumors. DAF-18 is largely dispensable in sperm-bearing hermaphrodites, potentially only ensuring a tighter control of oocyte maturation and ovulation (Brisbin et al., 2009; Suzuki and Han, 2006). Even in older sperm-depleted hermaphrodites or feminized mutants, and despite the resulting ongoing GSC proliferation, the loss of *daf-18(ø)* has no tumorous consequences since the continually produced oocytes are spontaneously ovulated and laid out through the vulva. In a background that prevents oocyte activation however, such as in the *oma-1; oma-2* double mutant (Detwiler et al., 2001), loss of the AMP activated protein kinase ortholog *aak-1*, a gene that is required for homeostatic downregulation of GSC proliferation like *daf-18*, led to the formation of a benign differentiated germline tumor since diakinesis-blocked oocytes were not laid and hyperaccumulated (Narbonne et al., 2017; Valet and Narbonne, 2022). Similar to *aak-1(*ø*); oma-1; oma-2* triple mutants, *daf-18(*ø*) oma-1; oma-2* triples grew oocyte tumors (Figure S6). Hence, we conclude that DAF-18 can cell non-autonomously prevent the formation of benign differentiated germline tumors through affecting contractility in the adjacent somatic tissue via AKT and RhoA.

## Discussion

Altogether, our data are consistent with a model in which DAF-18 acts in the Sp neck to promote its contractility in the absence of sperm, preventing the ovulation and wastage of unfertilized oocytes. This in turn allows for the homeostatic downregulation of GSC proliferation, also pausing oocyte production. In conditions where oocyte laying is prevented such as in the *oma-1; oma-2* background, DAF-18 activity become critical to prevent oocyte hyperaccumulation, or in other words, germline benign tumorigenesis. Homeostatic regulation of GSC proliferation could therefore arise as a downstream effect of unfertilized oocyte retention in the absence of sperm, resulting from RHO-1’s contribution to Sp neck contractility.

Even though Sp neck DAF-18 was sufficient for homeostatic downregulation of GSC proliferation, GSC proliferation was partially uncoupled from oocyte accumulation in feminized *daf-18(ø)* animal that were rescued specifically in the sheath cells (Figure 3E). As such, homeostatic regulation of GSC proliferation could involve a consequence of oocyte accumulation on the sheath cells that could be partially by-passed by *daf-18(+)* overexpression in this tissue. Indeed, as sheath MPK-1 activity promotes GSC proliferation (Robinson-Thiewes et al., 2021), while *daf-18(*ø) mutants have upregulated MPK-1 signaling (Nakdimon et al., 2012; Narbonne et al., 2017; Suzuki and Han, 2006) changes resulting from the lack of ovulation or oocyte accumulation could lead to the inhibition of MPK-1 signaling in the sheath cells and thereby downregulate GSC proliferation. Future investigations will be required to test this hypothesis.

Considering the previously established roles of DAF-18 in suppressing germline MPK-1 activity to ensure the proper timing of oocyte growth and maturation (Das and Arur, 2017; Narbonne et al., 2017; Suzuki and Han, 2006), it initially seemed reasonable to hypothesize that DAF-18 acted within the germline to suppress MPK-1 activation in the absence of sperm and block the final maturation of the oocyte by preventing it from releasing LIN-3 (Clandinin et al., 1998). This would have prevented the strong ovulatory contractions that lead to ovulation, and potentially be sufficient to trigger oocyte accumulation (Brisbin et al., 2009; Suzuki and Han, 2006). However, we found that restoring germline DAF-18 activity in *fog-1; daf-18(ø)* animals did not reinstate oocyte accumulation. Thus, our results establish that either *daf-18* functions non-autonomously in the soma to prevent oocyte MPK-1 activation in the absence of sperm, or that the spontaneous activation of MPK-1 within oocytes that happens in the absence of sperm and *daf-18* is insufficient to trigger ovulation. More analyses will be required to distinguish between these two possibilities.

Our findings showed that the reintroduction of DAF-18 in the non-pharyngeal muscles of *fog-1; daf-18(ø)* animals was sufficient to trigger oocyte accumulation and restore the downregulation of GSC proliferation. We realized that non-pharyngeal muscles encompass most tissues of the somatic gonad, including the sheath cells, Sp, and Ut, all of which are of myoepithelial nature. These tissues have been heavily implicated in the regulation of oocyte maturation and ovulation (Hiatt et al., 2009; McCarter et al., 1999; McGovern et al., 2007; Rose et al., 1997; Whitten and Miller, 2007), and thus were prime candidates for the critical role of *daf-18* towards ovulation control. Eventually, we narrowed down the minimal *daf-18* tissue requirements to the 8 cells that form the Sp neck (Castaneda et al., 2020). Remarkably, localized *daf-18* expression in the Sp neck also effectively quelled GSC proliferation, suggesting ovulation and GSC proliferation control are both linked to DAF-18 activity in the Sp neck. Should the suppression of MPK-1 activation within proximal oocytes by DAF-18 be non-autonomous, it could therefore potentially also stem from DAF-18’s role in the Sp neck.

We next delved into the mechanisms by which *daf-18*, non-autonomously from the Sp neck, ensures oocyte arrest and accumulation in the absence of sperm. We first assessed whether that function of DAF-18 was dependent on its lipid-phosphatase activity. There is considerable evidence from the mammalian literature that the G129E mutation selectively abrogates PTEN’s lipid phosphatase activity (Furnari et al., 1998; Han et al., 2000; Myers et al., 1997; Ramaswamy et al., 1999). In *C. elegans*, the corresponding DAF-18(G174E) mutation has been evaluated by four groups before us (Chen et al., 2022; Feng et al., 2018; Nakdimon et al., 2012; Wittes and Greenwald, 2022; Zheng et al., 2018). We note that two recent studies failed to confirm the specificity of the lipid phosphatase-defective allele, but that could be because the defects that they were investigating entirely depended on this activity (Chen et al., 2022; Wittes and Greenwald, 2022). On the other hand, Nakdimon *et al*. reported defects of *daf-18(ø)* that were rescued by the G174E allele, suggesting that this allele may indeed leave its protein phosphatase activity intact (Nakdimon et al., 2012). It may therefore be reasonably concluded that the control of ovulation and homeostatic regulation of GSC proliferation by DAF-18 depend on its lipid phosphatase activity.

Next, using video microscopy and a Ca^2+^ biosensor, we demonstrated that DAF-18 prevents Sp neck dilation, likely by increasing its contractility. We showed that DAF-18 accomplishes this independently from PLC-1/ITR-1/Ca^2+^ signaling despite the fact that DAF-18 should theoretically feed PIP_2_ into this pathway to promote contractility. Even though *itr-1(rf)* significantly prevented oocyte accumulation in feminized mutants, almost to the level of *daf-18(ø)*, and prevented *daf-18(ø)* from further decreasing oocyte accumulation (Figure 5F), we disregarded the hypothesis that *daf-18* was working mainly through the PLC-1/ITR-1/Ca^2+^ cascade because 1) basal Sp Ca^2+^ levels remained unaffected by *daf-18(ø)* in all tested backgrounds, and 2) the removal of *daf-18* still produced oocyte accumulation and/or GSC proliferation defects in both *plc-1(ø)* and *itr-1(rf)* mutants. Our results further established that PLC/ITR signaling is required for optimal GSC proliferation in sperm-bearing hermaphrodites (Figures 5E and 5G). Since *plc-1(ø)* and *itr-1(rf)* single mutants do not accumulate unfertilized oocytes (Figure 5D and 5E) but rather accumulate eggs in their Sp (Kovacevic et al., 2013), it raises the question of what exactly is the signal that triggers homeostatic downregulation of GSC proliferation. Is it really anovulated oocyte accumulation or could it simply result from the physical blockage of germ cell flux? If so, how could that blockage exactly promote GSC quiescence? Alternatively, could PLC/ITR signaling regulate GSC proliferation independently from their role in the Sp? Towards these quests, it will be important to investigate how *plc-1* and *itr-1* interact with IIS and MPK-1, the two established growth factor pathways that converge to determine GSC proliferation rates in *C. elegans* (Narbonne et al. 2017).

We found that the loss of DAF-18 increased Sp levels for its subtrate. This extra PIP_3_ is expected to activate the AKT-1/2 kinases (Paradis et al., 1999). Accordingly, we showed that AKT-1/2 are then required to inappropriately promote ovulation in *daf-18(ø)* mutants and make it happen even in the absence of stimulation by sperm. This occurs independently from DAF-16/FOXO, but instead likely involves the direct phosphorylation of RHO-1 on a perfectly conserved site by AKT-1/2. We show that such phosphorylation would inhibit RHO-1 activity, and results in a reduced contractile response to normal Ca^2+^ influxes in the Sp neck, facilitating its dilation and ovulation. Interestingly, it was reported in mice that a cardiomyocyte-specific deletion of PTEN caused a dramatic decrease in cardiac contractility that depended on PI3Kγ (Crackower et al., 2002). The positive regulation of contractility by PTEN via the regulation of PIP_3_ levels may therefore be conserved.

Given the high level of evolutionnary conservation of the signaling proteins that we investigated here in *C. elegans*, and particularly of the new putative AKT-1/2 phosphorylation site on RHO-1 that was uncovered, it appears likely that AKT could be influencing RhoA activity, potentially through direct phosphorylation, in a variety of contractile or secretory tissues in other organisms, including humans, and modulate how they respond to Ca^2+^ influxes. If these tissues were to serve as sensors or gauges that monitor tissue homeostasis in a manner homologous to how the Sp neck influences germline homeostasis in *C. elegans*, this cascade could equally impinge on stem cell regulation and prevent benign tumorigenesis in other systems. PTEN could therefore act through similar cell non-autonomous mechanisms in human to prevent the formation of benign differentiated tumors in multiple tissues where stem cell proliferation rates may be coupled to the need for their differentiated cellular progeny. As such, the systemic reduction of PTEN activity that characterizes PHTS patients could therefore predispose individuals to the formation of hamartomas, benign differentiated tumors, through impairing homeostatic regulation of stem cell proliferation (Valet and Narbonne, 2022).

## Materials and Methods

### C. elegans genetics

Animals were maintained at 15LJ on standard NGM plates and fed *E. coli* bacteria of the strain OP50, unless otherwise indicated (Brenner, 1974). The Bristol isolate (*N2*) was used as wild type throughout. All alleles, deficiencies and transgenes used are listed in Table S1.

### Plasmids and transgenics

We used the Gibson (Gibson et al., 2009) method for assembling all plasmids. The source DNA and primers that were used to generate all plasmids, as well as their microinjection concentrations, are listed in Table S2.

Extra-chromosomal arrays were generated by standard germline microinjections at a total concentration of 200 ng/μL, using pKSII as filler DNA and pCFJ104[*Pmyo-3::mCherry*] (5 ng/μL) or pMR352[*Pmyo-2::GFP*] (Narbonne and Roy, 2009) (50 ng/μL) as co-injection markers (Frokjaer-Jensen et al., 2008; Mello et al., 1991). For *Punc-54*, we used the fragment driving enhanced expression previously coined “*PEunc-54*” (Masse et al., 2005). To rescue DAF-18 specifically in the germline, we used CRISPR/Cas9 to insert a single copy of a *Pmex-5::GFP::DAF-18(+)* + *unc-119(+)* fragment at ttTi5605 (LG II, +0.77 M.U.) into *unc-119(ed3)* mutants, using pPOM4 (Table S2) and pDD122 (Dickinson et al., 2013). A single line (*narSi5*) was obtained after injection of > 80 animals. To generate the *daf-18(G174E), rho-1(S73A)* and *rho-1(S73D)* variants, we used the *dpy-10* co-CRISPR strategy with the Paix *et al*. protocol (Dickinson et al., 2015; Paix et al., 2015). We grew P0s on *cku-80(RNAi)* to favor all CRISPR/Cas9 genomic editions (Ward, 2015). The sgRNAs and repair template sequences used are listed in Table S3.

### Oocyte counts

Late-L4 stage hermaphrodites were transferred from 15LJ to a new plate at 25LJ and after 3 days, F1 late-L4s hermaphrodites were synchronized by picking them to a new plate at 25LJ based on vulva development (Seydoux et al., 1993). They were grown for an additional 24 hours (Narbonne et al., 2015). This procedure allowed to inactivate the temperature sensitive *fog-1(q253)* allele throughout F1 larval development to prevent sperm formation (Barton and Kimble, 1990). Resulting feminized day 1 adults (A1) were harvested, paralyzed with 0.1% Tetramisole (Sigma, L9756) and mounted onto M9 + 3% agarose pads. The number of oocytes per gonad arm, and their compaction status, were determined by differential interference contrast (DIC) examination.

### Germline mitotic index

Progenitor zone mitotic indexes were evaluated as previously described (Crittenden et al., 2006; Hubbard and Schedl, 2019; Narbonne et al., 2015, 2017; Robinson-Thiewes et al., 2021). Synchronized A1 hermaphrodites were generated as above and their gonads were dissected and stained as previously described (Robinson-Thiewes et al., 2021). Briefly, hermaphrodites were transferred into an 8 µL drop of 1X PBS on a microscope slide cover glass and quickly dissected using a 25G surgical needle tip. The cover glass was then flipped onto a poly-L-lysine coated slide and submitted to a freeze-crack procedure. Samples were fixed in -20LJ methanol for 1 minute and postfixed in a 3.7% paraformaldehyde solution (3.7% paraformaldehyde [PFA], 1X PBS, 0.08 M HEPES, 1,6 mM MgSO4 et 0,8 mM EGTA) for 30 minutes. Using 1ml PBST each time, (PBS + 0.1% Tween 20) slides were rinsed twice followed by a 10 minutes incubation in PBST. Samples were then covered with 300 µL blocking solution (PBST + 3% BSA) for 1h at room temperature. Samples were stained using primary rabbit anti-WAPL-1 (1:500, Novus Biologicals #49300002) to mark GSCs along with their proliferating progeny (Kocsisova et al., 2018), and mouse anti-phospho[ser10]-histone H3 antibodies (1:250, Cell Signaling #9706) to mark G2/M-phase nuclei, and counter-stained with 0.7 µg/mL 4’6-diamidino-2-phenylindole (DAPI) to highlight all nuclei. Progenitor zone nuclei counting in 3 dimensions was partially automated using an ImageJ plugin developed by Dr Jane Hubbard’s laboratory (Korta et al., 2012).

### Image acquisition and processing

For Figures 1C, 2A, 3B, 4A, 6B, 6D, 7E, 7H, S2, S3B, S4A, and S6, DIC and/or epifluorescence z-stacks were acquired every micron using a Plan-Apochromat 20x dry objective (NA 0.8) mounted on an inverted Zeiss Axio Observer.Z1. Micrographs were stitched using the Zen software (2.6 Blue Edition) and animals were straightened for ease of visualization using ImageJ. Epifluorescence signals were overlaid to the DIC micrographs using ImageJ.

For the high-resolution confocal fluorescence acquisitions in Figure 1A, 5C and S3C, A1 hermaphrodites were anaesthetized and mounted for imaging as above. Coverslips were sealed with VALAP (1:1:1 vaseline, lanolin, paraffin), and 0.35 μm z-step stacks were acquired using a Leica SP8 point scanning confocal microscope with an HC PL APO CS2 63x/1.30 numerical aperture oil objective.

For Figures 5A, and Videos S1-3, z-stack DIC, epifluorescence micrographs and video were acquired every micron using a Plan-Apochromat 40x/1.4 oil objective mounted on an inverted Zeiss Axio Observer.Z1. Epifluorescence signals were overlaid to the DIC micrographs using ImageJ. All DIC micrographs show single focal planes, and where applicable, overlaid fluorescent micrographs also show the corresponding single focal plane.

For MI evaluations (Figures 1E, 2C, 3D, and 4C), representative maximal projections from z-stacks epifluorescence micrographs of the distal gonad acquired every micron for Alexa 488, Alexa 546 and DAPI using a Plan-Apochromat 40x/1.4 oil objective mounted on an inverted Zeiss Axio Observer.Z1 are shown.

### DAF-18::mNG quantification

Synchronized A1 hermaphrodites were generated, paralysed and mounted on a M9 + 3% agarose pad as above. High-resolution images of DAF-18::mNG were acquired using a Leica SP8 point-scanning confocal microscope with an HC PL APO CS2 63x/1.30 numerical aperture oil objective, using z-stacks at 0.35 μm intervals. Z-stacks were merged into single images using maximum intensity projection in ImageJ. A 10-pixel-wide segmented line was manually drawn along the oocyte membranes separating the -1 and -2 oocytes to measure their average fluorescence intensity.

### Dauer formation assays

Dauer formation was scored as described in previous reports (Paradis and Ruvkun, 1998). Briefly, for Figure S1B, bleached batches of eggs were allowed to hatch in the absence of food at 15LJ for 36 hours and synchronized L1s were plated at 25°C to induce dauer entry. Dauer formation rate was evaluated as the number of dauers divided by the total number of larvae after 96 hours. For Figure S1C and S1D however, hermaphrodites were picked together with males at the L4 stage and allowed to mate for 24 hours. Hermaphrodites were then singled onto new plates at 25°C and dauer formation rates in the crossed F1 progeny, or F2 progeny from crossed F1, was evaluated.

### Ca^2+^ imaging and quantification

Intracellular Sp Ca^2+^ was evaluated in A1 hermaphrodites using the *Pfln-1::GCaMP3 in vivo* sensor (a kind gift from Erin J. Cram) (Kovacevic et al., 2013). Fluorescence intensity profiles were obtained using confocal microscopy and processed using ImageJ as previously described (Kovacevic et al., 2013). Briefly, animals were paralyzed in a 0.1% tetramisole (Sigma, L9756) M9 solution and mounted on a 3% agarose pad. Coverslips were sealed with VALAP (1:1:1 vaseline, lanolin, paraffin), and 0.35 μm step z-stacks were acquired using a Leica SP8 point scanning confocal microscope with an HC PL APO CS2 63x/1.30 numerical aperture oil objective. A maximal z-projection was performed to obtain display micrographs. A segmented line was used to define a polygonal region of interest (ROI) corresponding to the Sp. The mean fluorescence intensity was extracted for each ROI and background subtracted.

### PIP_3_ staining and quantification

A1 hermaphrodites were harvested and their gonads were dissected and stained as described for germline mitotic index evaluations above (Robinson-Thiewes et al., 2021). Primary mouse monoclonal anti-PIP_3_ antibodies (1:100, Echelon Z-P345) and rabbit anti-HIM-3 (a kind gift from M. Zetka), and secondary A488-conjugated goat anti-mouse or A546-conjugated goat anti-rabbit antibodies (both at 1:500, Invitrogen Cat# A-32731 and A-11035, respectively) were used. DAPI was used as a counterstain. For Figure 6A, micrographs were acquired every micron using a 60x oil immersion objective mounted on a DeltaVision microscope, maximally-projected, stitched, straightened and thresholded using ImageJ. For Figure 6B, z-stack DIC and fluorescence micrographs were acquired every micron using a Plan-Apochromat 40x/1.4 oil objective mounted on an inverted Zeiss Axio Observer.Z1. The micrographs were then transferred to Imaris 9.2.1, and based on the green channel (A488), 3D models were individually thresholded to best fit the spermatheca contour. For PIP_3_ quantification, the GFP object was selected, and using the edit tool, its mean intensity was pulled from the statistics tab.

### Endomitotic oocyte laying quantification

Synchronized A1 hermaphrodites were generated as above and singled to a new plate at 25°C for an additional 24 hours. Adults were removed and the number of endomitotic oocytes laid during this 24-hour period was determined under a dissecting microscope.

### Developmental Milestone Analysis

Developmental milestone analysis was scored as described in previous reports (Woodruff et al., 2019). Briefly, a large population of *C. elegans* was synchronized by bleaching as described above for dauer formation analyses. L1s were placed on seeded plates at the 25°C temperature and monitored every 24 hours. Developmental progression was tracked by counting the number of larvae that had reached or developed beyond the L4 stage. For analysis, animals were plotted by their developmental status (“0” = yet to reach milestone; “1” = reached milestone).

### In silico protein modeling and docking predictions

The predicted ternary structures of AKT-1 and RHO-1 were generated using SWISS-MODEL (Guex et al., 2009; Waterhouse et al., 2018). Their amino acid sequences were aligned against the Protein Data Bank (PDB), and the crystal structures 1H10 and 3CQW were selected as templates to fold AKT-1 and RHO-1, respectively. We used the High Ambiguity Driven Protein-Protein DOCKing (HADDOCK) 2.4 (de Vries et al., 2010) platform to predict the physical interaction between the AKT-1 kinase domain and RHO-1’s S73 site, and to obtain docking scores predictive of the likelihood of such an interaction. The HDOCK platform was also used to confirm favorable docking scores.

### Statistical analysis

Normality was verified using the Shapiro-Wilk test. Variance was verified with the F test for pairwise comparisons and using the Brown-Forsythe test for datasets comprising more than two groups. Tests used are indicated in the figure legends and were chosen according to the following criteria. For pairwise comparisons with normal distributions and equal variances, we used the unpaired two-tailed t-test. For non-gaussian distributions, the Mann-Whitney test was instead used. When the distribution was normal, but the variance was unequal, we used the Welch’s t-test. For multi-group comparisons that all had normal distributions and equal variances, we used the one-way ANOVA with Tukey’s multiple comparisons. For non-gaussian distributions, we used the Kruskal-Wallis with Dunn’s multiple comparisons. When the distribution was normal, but the variances were unequal, we used the Brown-Forsythe and Welch’s ANOVA with Dunnett’s T3 multiple comparisons test. Graphs were generated using GraphPad Prism 8. Asterisks indicate statistical significance (***: P<0.001; **: P<0.01; *: P<0.05) to all other samples unless otherwise specified. Statistical details, including all sample sizes, are indicated in the figure legends.

## Acknowledgements

We thank Jean-Claude Labbé for constructive comments and edits on the manuscript; Erin J. Cram, Monique Zetka and Florence Solari for reagents and strains; the *Caenorhabiditis* Genetics Center (CGC) and WormBase for their essential roles in *C. elegans* research. The work in the Simard laboratory was supported by The Canadian Institutes of Health Research (CIHR). Research in the Narbonne laboratory is funded by grants from the NSERC (RGPIN-2019-06863, RGPAS-2019-00017, DGECR-2019-00326), the CIHR (PJT-169138), and a Research Chair from the *Fondation Marcel et Rolande Gosselin* to PN. PN is a junior 2 FRQS bursary scholar (310643).

## Author Contributions

JD: Study design, experimentation, data analysis, manuscript drafting.

AMC: Helped generating UTR507 and contributed Figures 7B-D, S5A, video S4.

OG: pOG1 design and construction, corresponding lines derivation and preliminary analyses; contributed to Figure 3B.

VR: Key preliminary analyses towards Figures 2A and S1.

BD: UTR29 derivation and preliminary analyses towards Figure S6. POM: pPOM4 design and construction.

MJS: Supervision and manuscript editing.

PN: Study design, experimentation, supervision, manuscript drafting and editing.

---

**Table S1.**
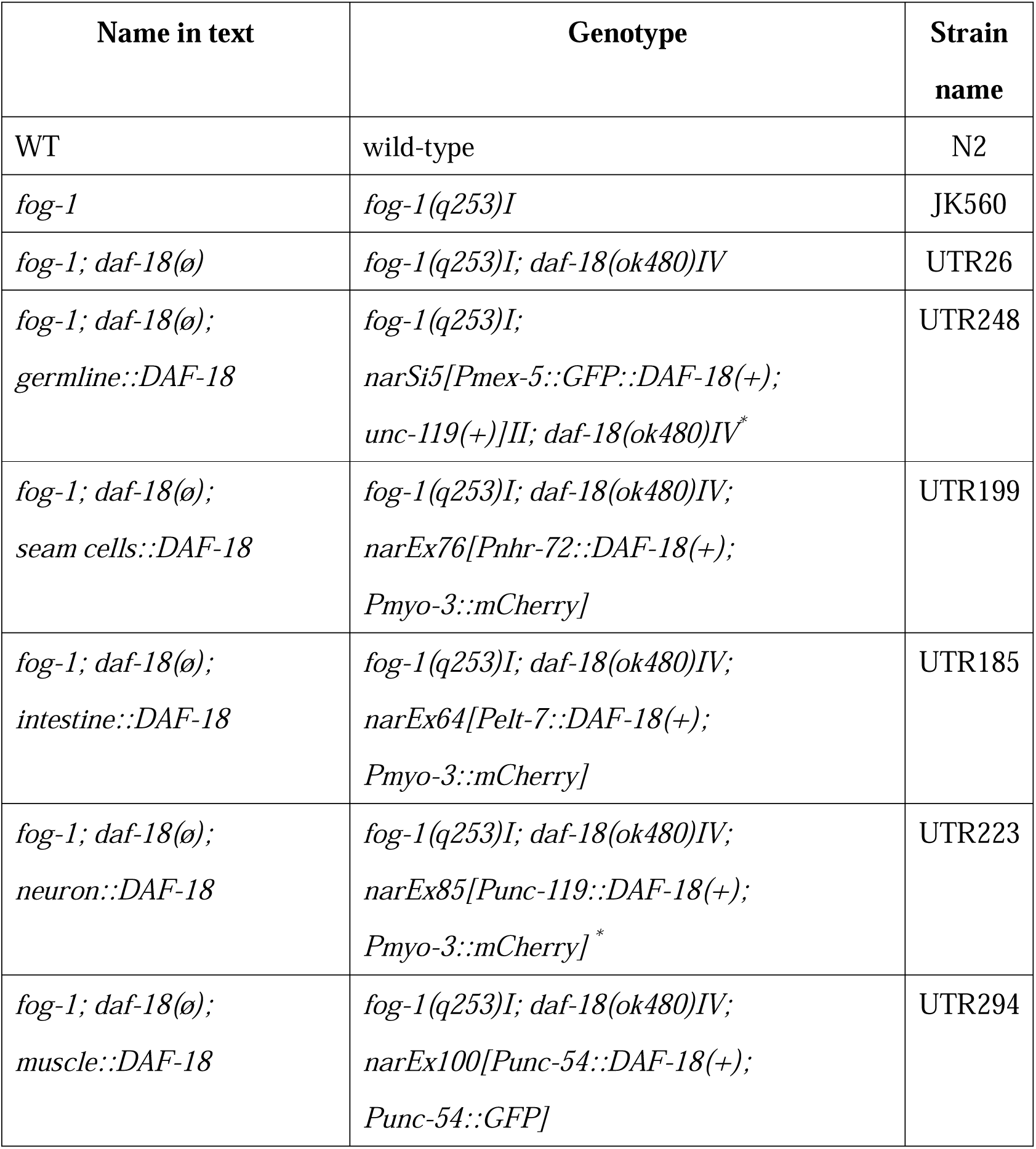
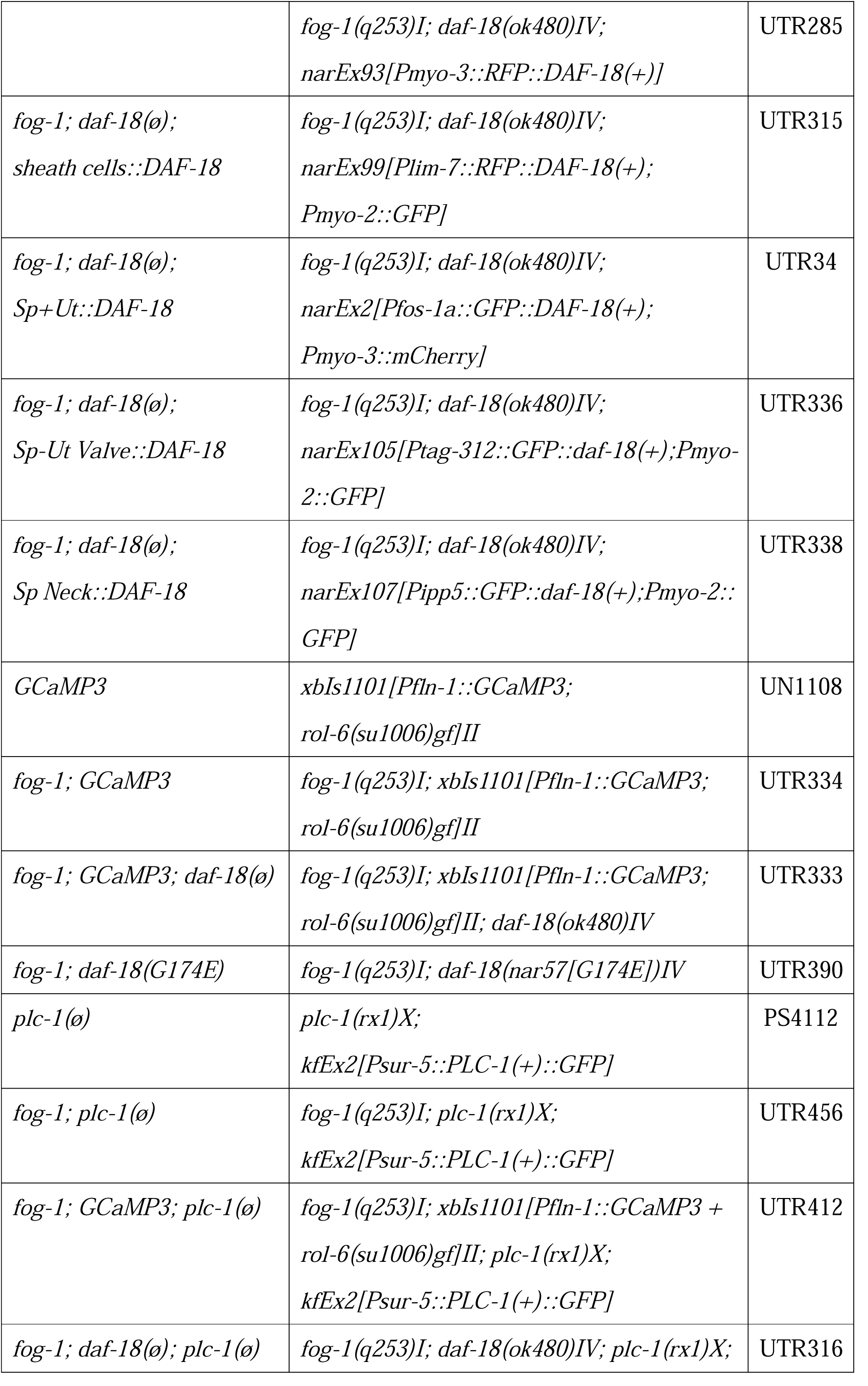
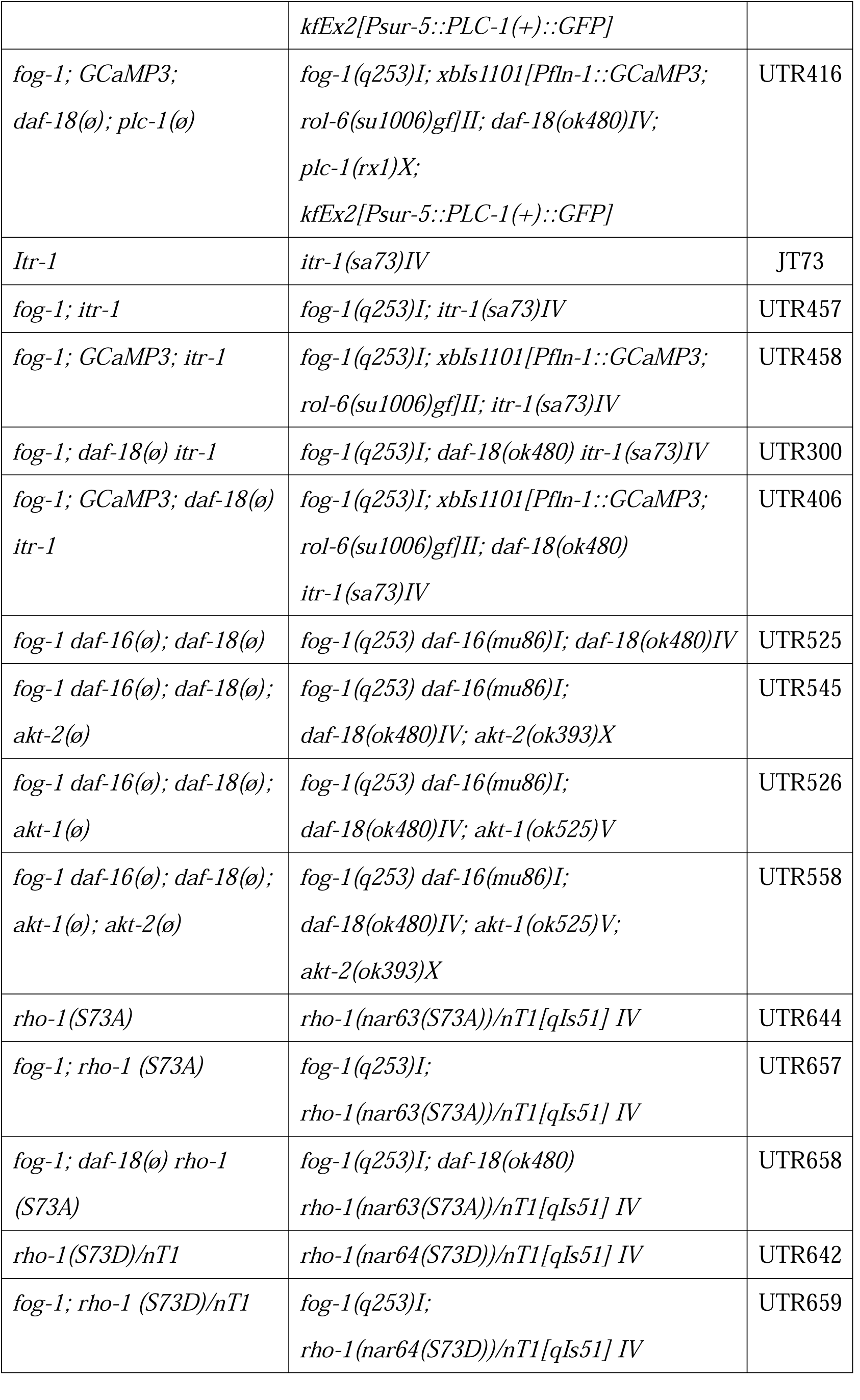
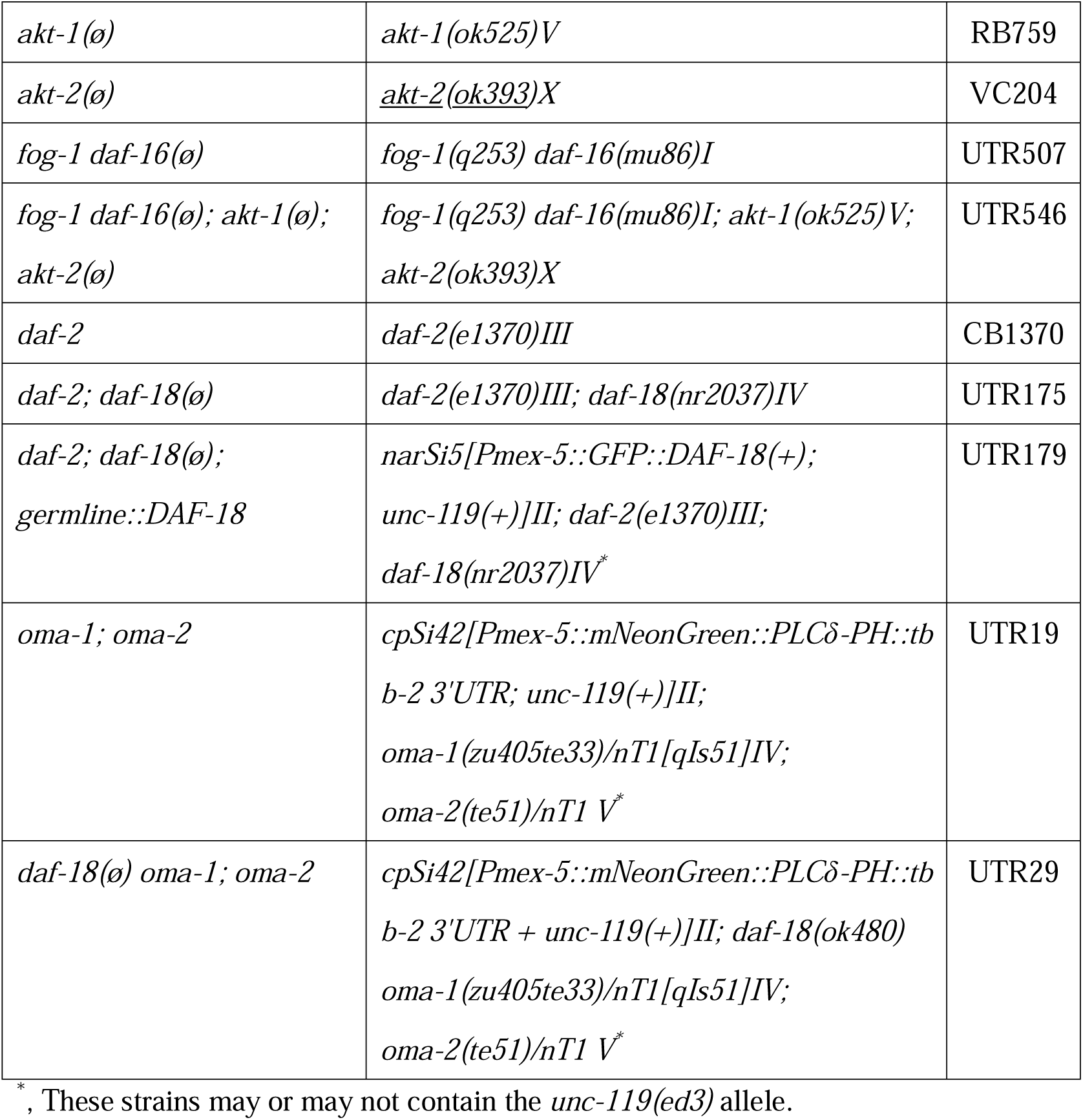
Strains, alleles, transgenes and rearrangements used.

**Table S2.** Plasmid design.

| Plasmid name | Description | Injection concentration | Reference |
| --- | --- | --- | --- |
| pKSII | Empty cloning vector | 100ng/μL | N/A |
| pPOM4 | *Pmex-5::GFP::DAF-18(+)* for single-copy insertion on LGII | 50ng/μL | This work |
| pFS2 | *Pelt-7::DAF-18(+)* | 50ng/μL | Masse et al., 2005 |
| pFS4 | *Punc-119::DAF-18(+)* | 50ng/μL | Masse et al., 2005 |
| pFS5 | *Punc-54::DAF-18(+)* | 50ng/μL | Masse et al., 2005 |
| pFS6 | *Pnhr-72::DAF-18(+)* | 50ng/μL | Masse et al., 2005 |
| pJC3 | *Pmyo-3::RFP::DAF-18(+)* | 50ng/μL | This work |
| pFS9 | *Punc-54::GFP* | 50ng/μL | Masse et al., 2005 |
| pOG1 | *Pfos-1a::GFP::DAF-18(+)* | 50ng/μL | This work |
| pJC4 | *Plim-7::RFP::DAF-18(+)* | 50ng/μL | This work |
| pJC10 | *Pipp-5::GFP::DAF-18(+)* | 50ng/μL | This work |
| pJC14 | *Prsef-1::GFP::DAF-18(+)* | 50ng/μL | This work |
| pCFJ104 | *Pmyo-3::mCherry* | 5 ng/μL | Frøkjær-Jensen et al., 2008 |
| pMR352 | *Pmyo-2::GFP* | 50ng/μL | Narbonne and Roy, 2009 |

**Table S3.**
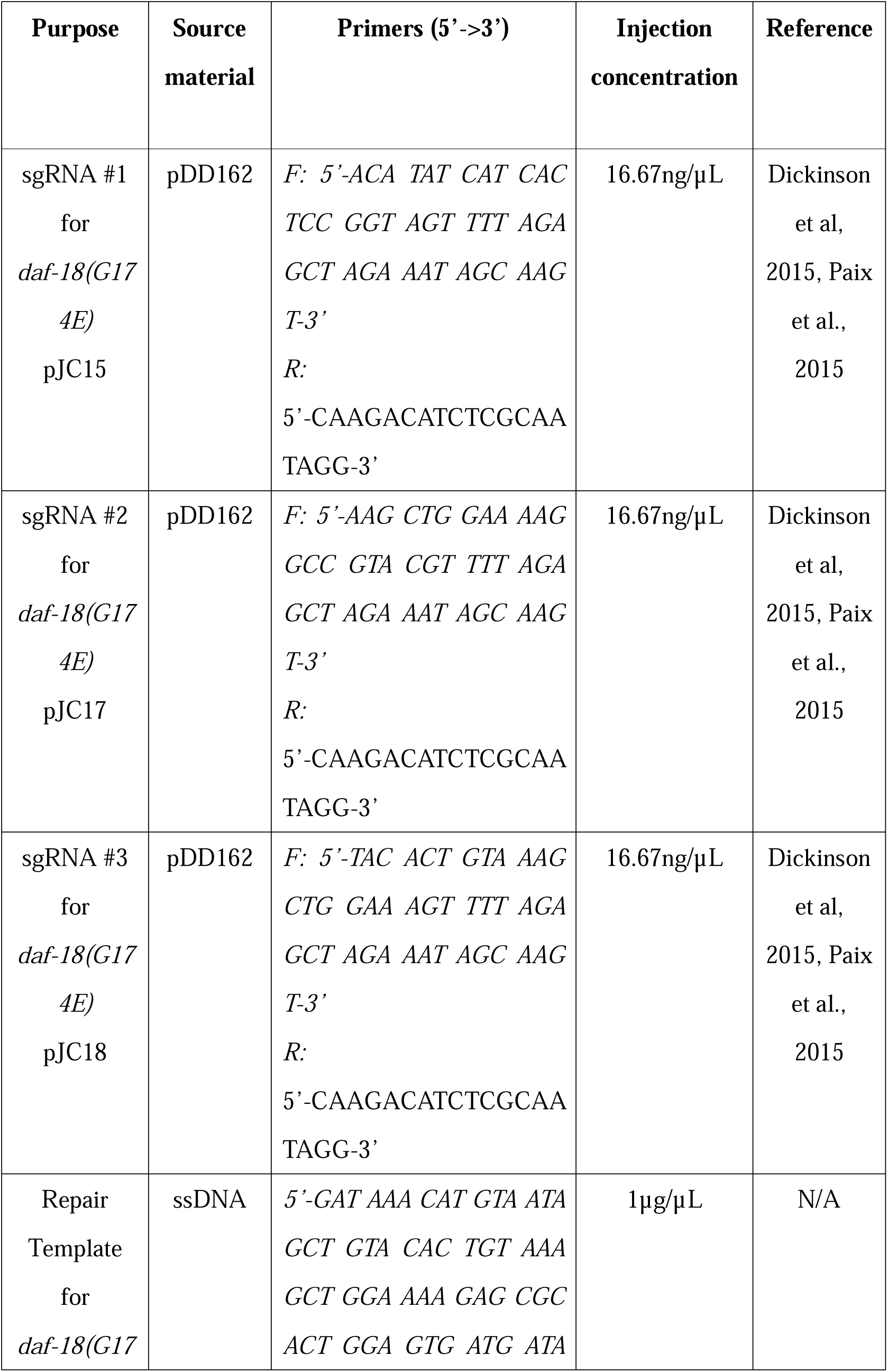
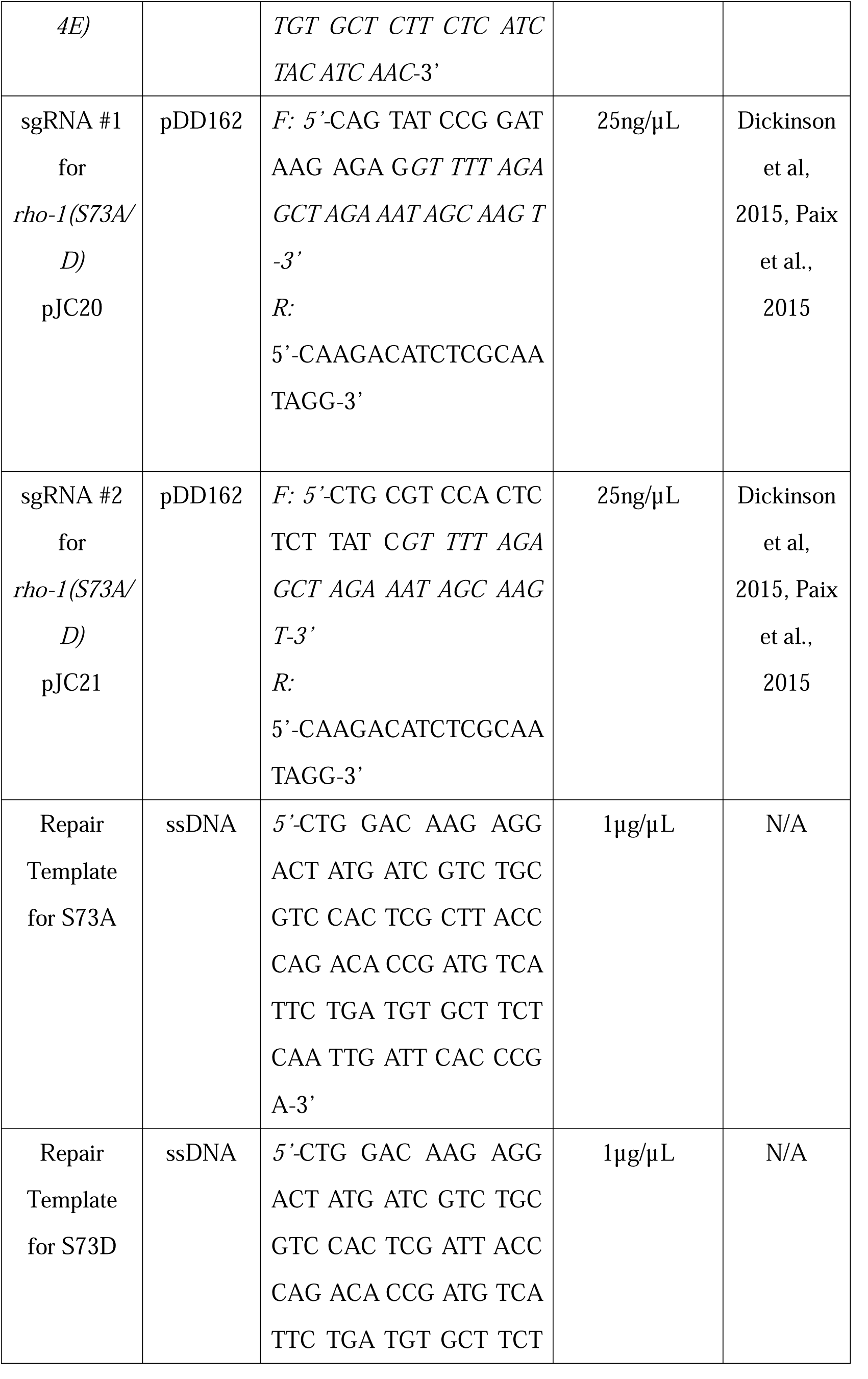
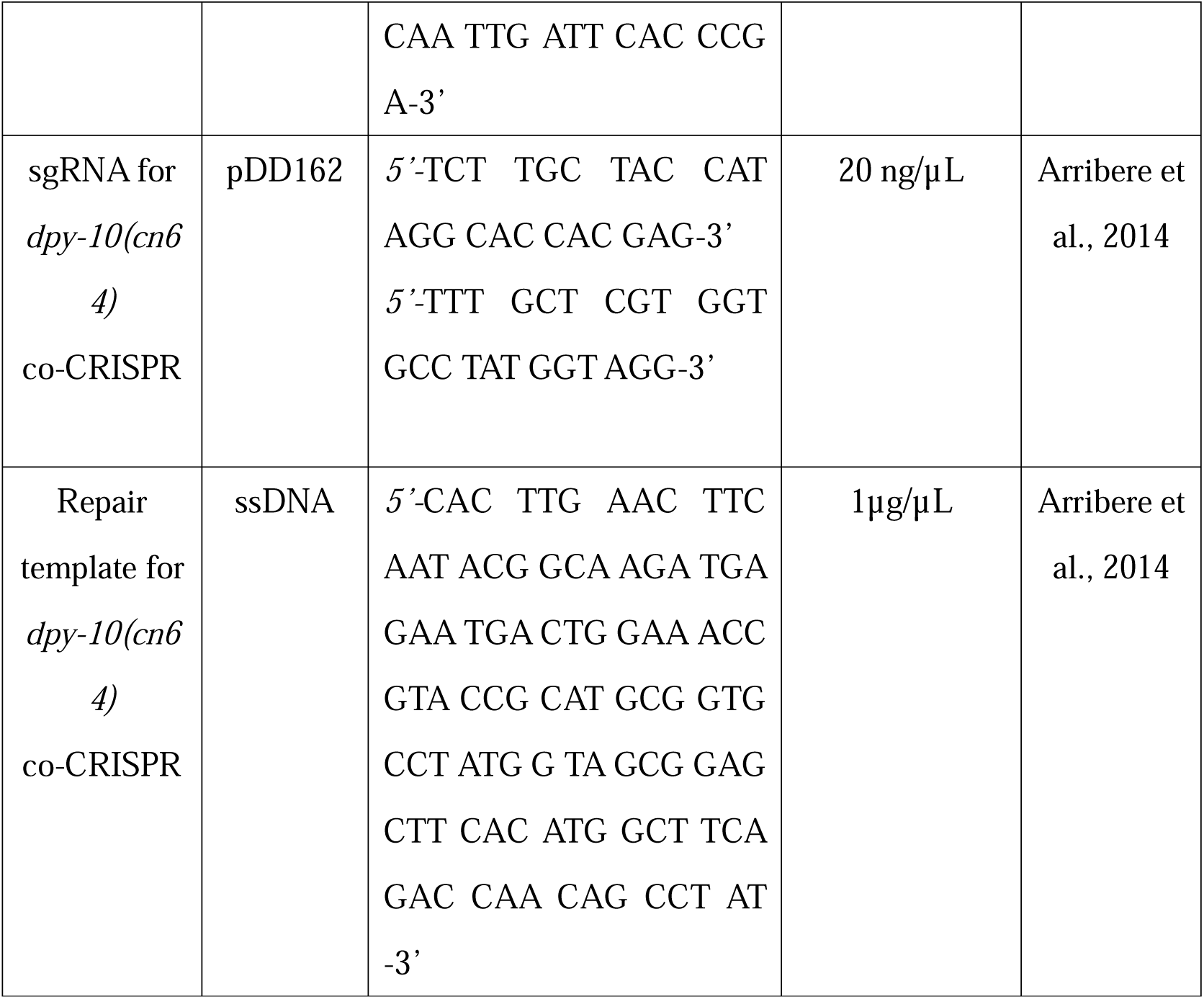
CRISPR design.

