## Supplementary figures and images for "DAF-18/PTEN prevents oocyte wastage in spermless *C. elegans* hermaphrodites by blocking spermatheca neck dilation through RHO-1/RhoA disinhibition"

### Figure S1

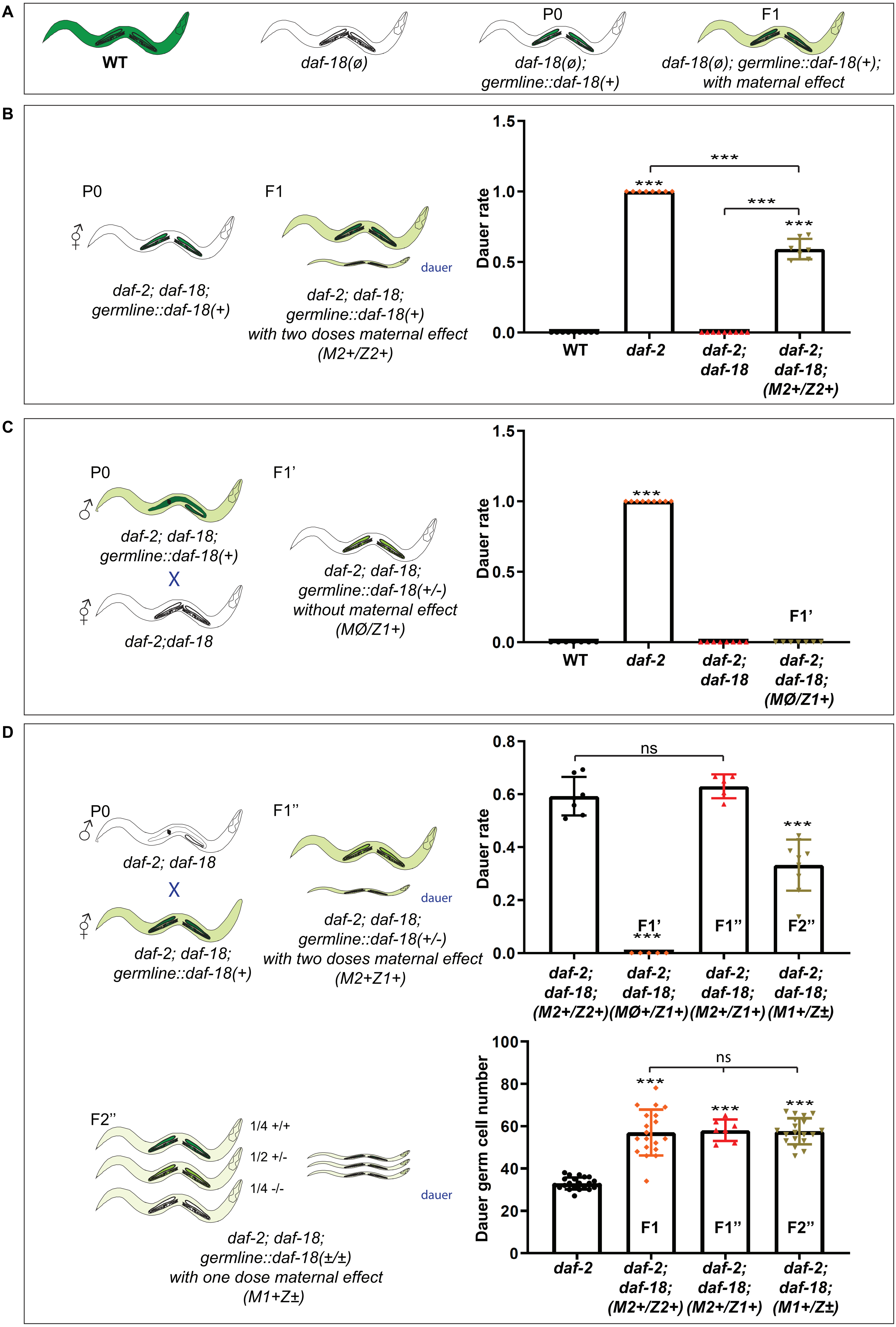

### Figure S2

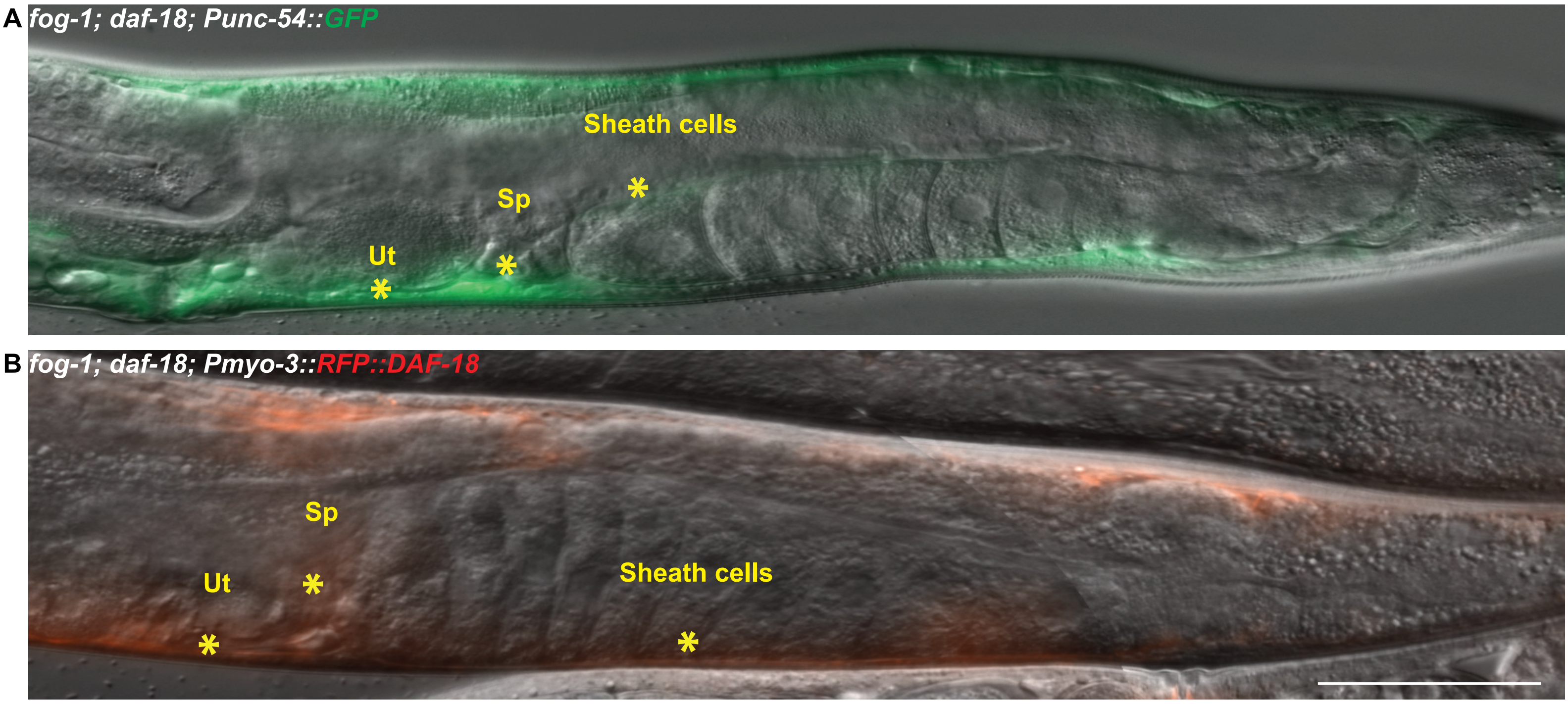

### Figure S3

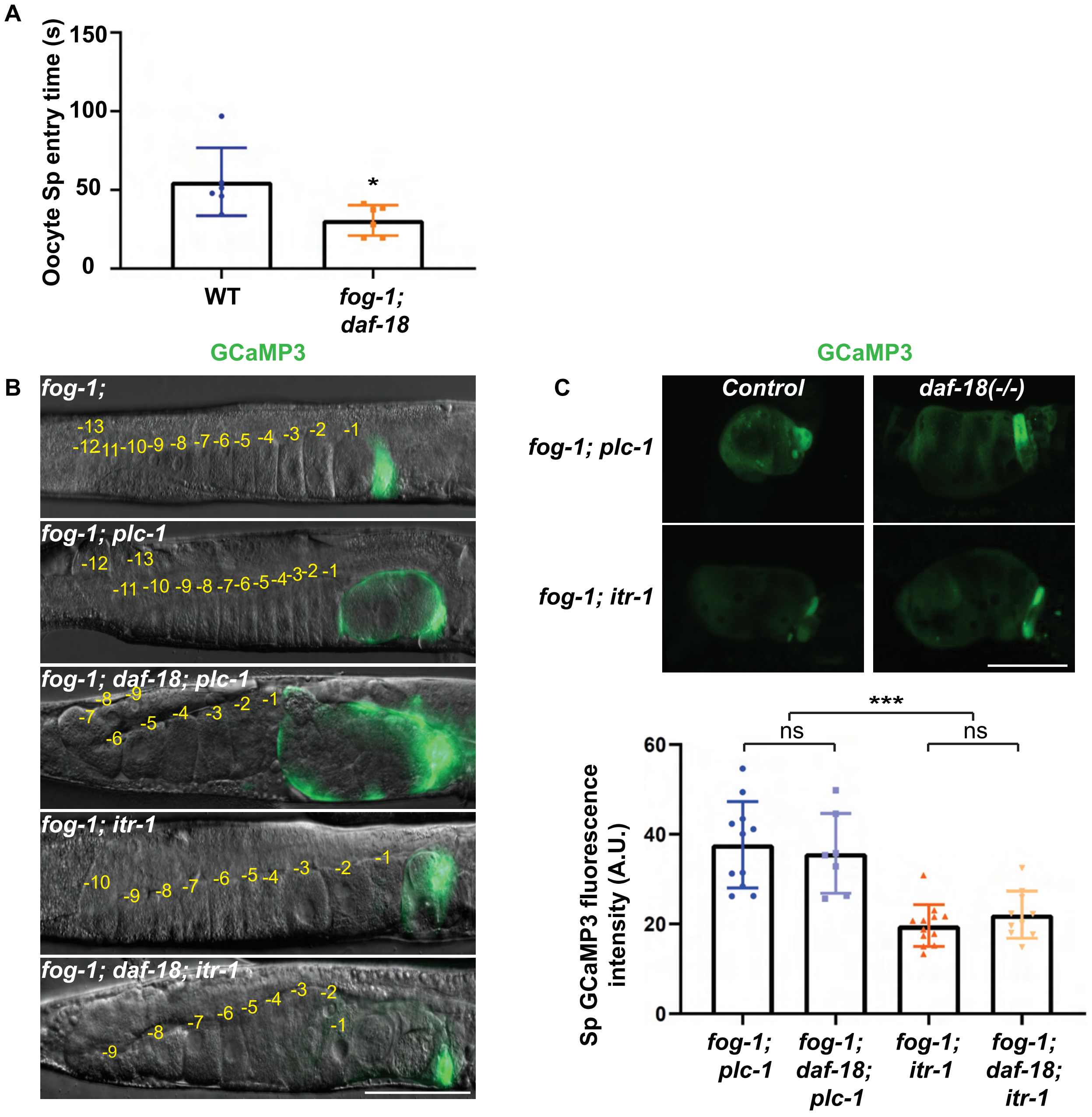

### Figure S4

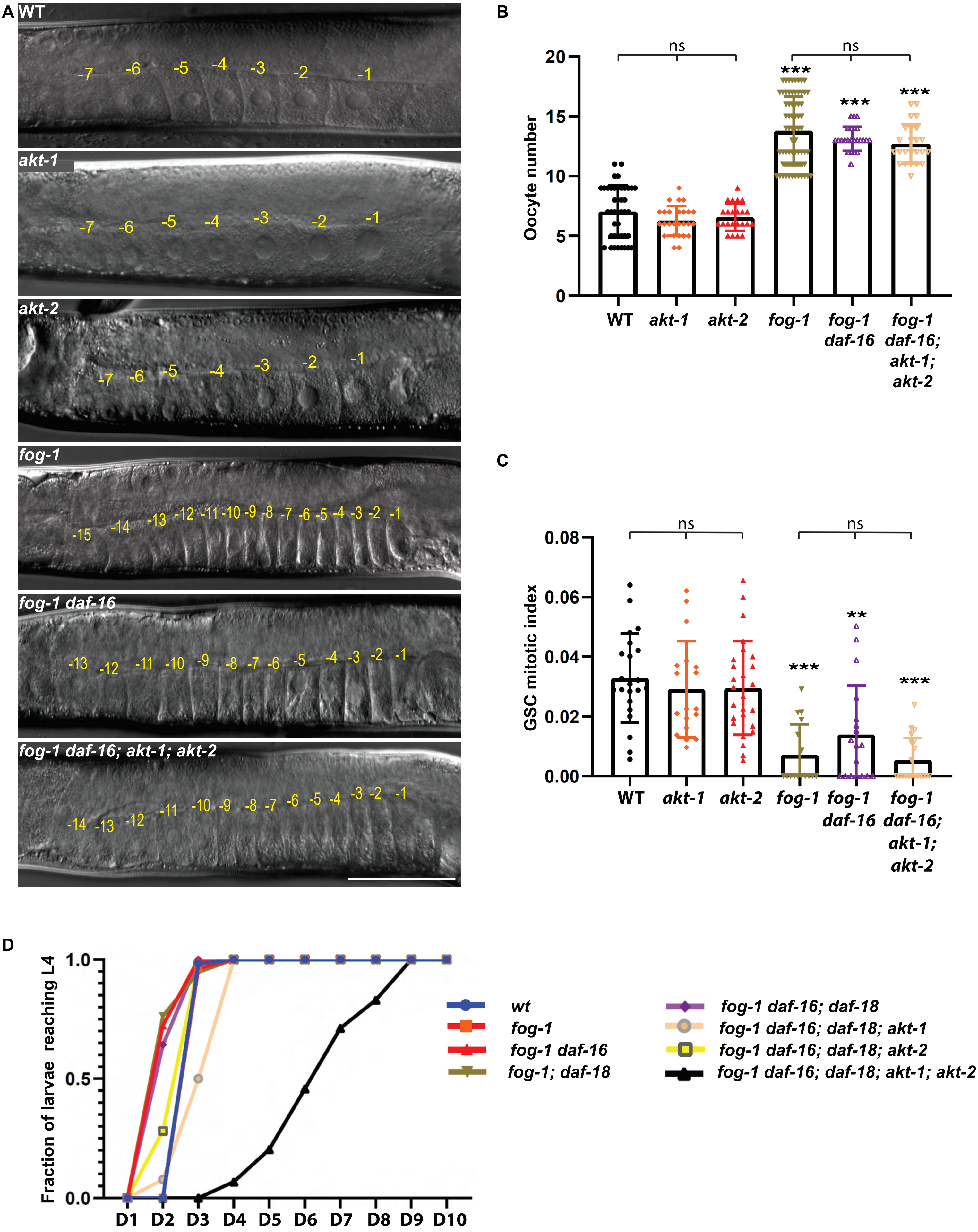

### Figure S5

A

| PROTEIN | AKT-1 HDOCK SCORE |
| --- | --- |
| RHO-1 | -244.11 |
| DAF-16 | -262.87 |

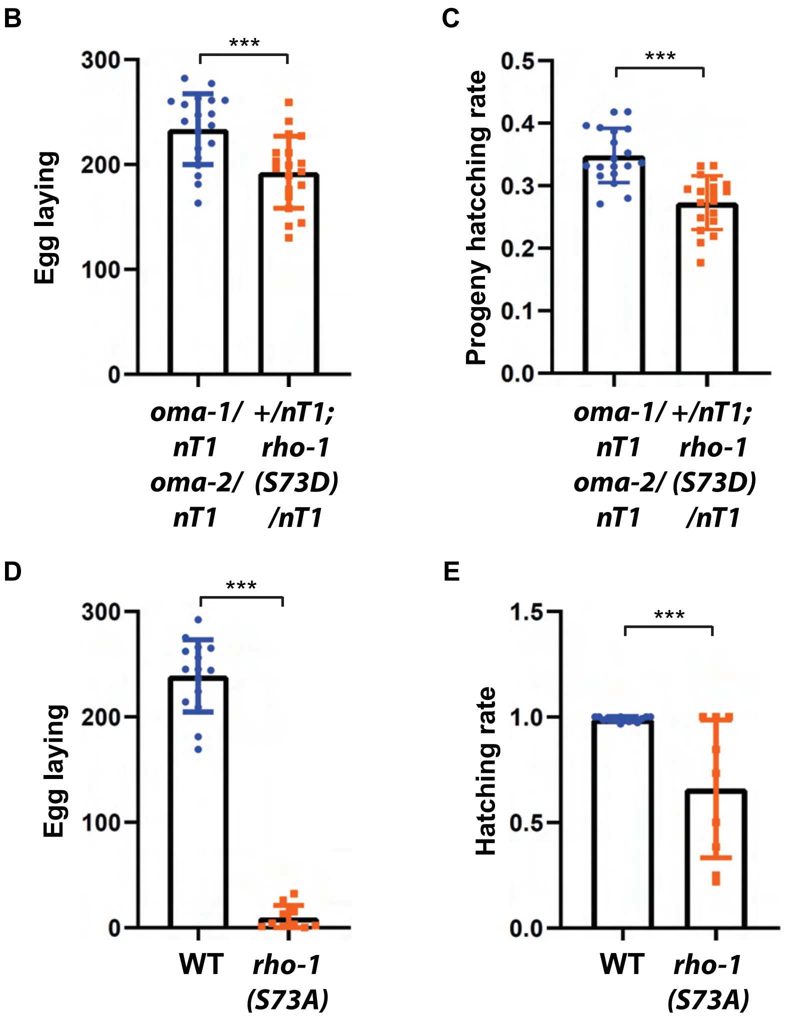

### Figure S6

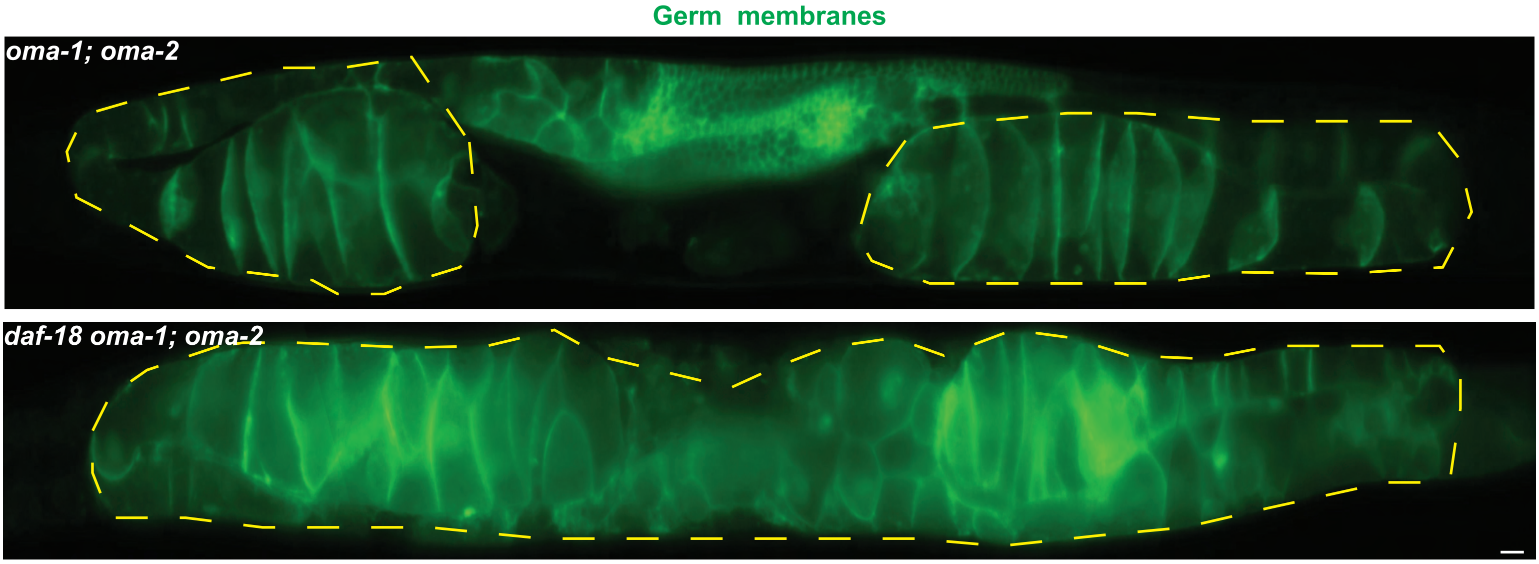
